# VPAC2 receptor signaling promotes pancreatic cancer cell growth and decreases the immunogenicity of the tumor microenvironment

**DOI:** 10.1101/2024.01.16.575872

**Authors:** Tenzin Passang, Shuhua Wang, Hanwen Zhang, Fanyuan Zeng, Po-Chih Hsu, Wenxi Wang, Jian Ming Li, Yuan Liu, Sruthi Ravindranathan, Gregory B. Lesinski, Edmund K. Waller

## Abstract

Identifying mechanisms underlying tumor growth and immune resistance is needed to treat pancreatic ductal adenocarcinoma (PDAC) effectively. The complexity of the tumor microenvironment (TME) suggests that the crosstalk between cells in the TME could drive drug resistance and relapse in PDAC. We have previously determined that vasoactive intestinal peptide (VIP) is overexpressed in PDAC and that VIP receptors expressed on T cells are a targetable pathway that sensitizes PDAC to anti-PD1 therapy. In this study, we show that pancreatic cancer cells engage in autocrine signaling of VIP through VIP-receptor 2 (VPAC2), and that high co-expression of VIP with VPAC2 leads to reduced relapse-free survival in PDAC patients. Mechanistically, we identified piwi-like RNA-mediated gene silencing2 (Piwil2) as a tumor-cell intrinsic protein downstream of VPAC2 that regulates cancer cell growth. In addition, we discovered TGFβ-1 as a potential tumor-extrinsic inhibitor of T cell function induced by VPAC2 signaling. *In vivo*, knock out and knockdown of VPAC2 on PDAC cells led to reduced tumor growth rate and increased sensitivity to anti-PD-1 therapy in various mouse models of PDAC that were T-cell dependent. Overall, these findings emphasize the implications of VIP/VPAC2 signaling in the PDAC tumor microenvironment and further support the rationale for developing VPAC2-specific antagonists.

**Significance:** The autocrine VIP signaling via VPAC2 promotes cancer cell growth and dampens T cell function in pancreatic ductal adenocarcinoma and thus represents a potential therapeutic target in PDAC.

## Introduction

Pancreatic Ductal Adenocarcinoma (PDAC) is a highly lethal disease of the pancreas with a dismal 5-year overall survival rate of only 12% (1). By 2040, it is projected to become the second leading cause of cancer-related deaths in the United States (2). Approximately 80% of patients are diagnosed with non-resectable disease and are treated with standard of care, including radiation, chemotherapies, and targeted therapies, albeit with limited response (3–5). PDAC has a highly desmoplastic and immunosuppressive tumor microenvironment (TME) with a paucity of effector T cells that limits the effectiveness of anti-CTLA-4 and anti-PD1 monotherapies (6–8), further emphasizing the need to understand the biological underpinnings of PDAC that lead to immune resistance. Our group previously showed that PDAC overexpression of vasoactive intestinal polypeptide (VIP) enables cancers to evade immune surveillance (9).

VIP is a 28-amino acid peptide secreted by the enteric nervous system with known effects on peripheral vasodilation, cardiac contractility, and gut peristalsis (10). The biological actions of VIP are mediated by a family of G protein-coupled receptors, namely, VPAC1 and VPAC2 (11,12). In addition to its previously known functions, growing research indicates that VIP has anti-inflammatory effects on T cells, macrophages, and plasmacytoid dendritic cells (13–17). Our published data demonstrate that VIP expression by pancreatic cancer cells leads to paracrine signaling on T cells via VIP receptors, resulting in T cell suppression and resistance to anti-PD-1 therapy (9). We observed that pancreatic cancer cells also express VPAC1 and VPAC2 and that absence of VPAC2 on cancer cells results in delayed tumor growth rate *in vivo*, suggesting a potential autocrine function of VIP via VPAC2 (9).

A small body of literature indicates the growth-promoting properties of VIP in pancreatic cancer cell lines, which can be reversed by VIP-receptor antagonism (18–21). However, little is known about VIP receptors’ intracellular signaling and downstream mechanism. Moreover, there is a lack of clear understanding of the significance of VIP-receptor-expressing pancreatic cancer cells in the TME. In particular, limited studies have used clinically relevant orthotopic models to delineate the biological underpinnings of VIP receptors in PDAC.

This current work extends the understanding of how VIP receptors, mainly VPAC2, regulate tumor growth and modulate the TME. In PDAC human patients, higher VIP and VPAC2 expression correlate with worse relapse-free survival compared to patients with low VIP and VPAC2 expression. Mechanistically, VPAC2 deletion reduces *in vitro* colony formation by decreasing stem-cell relating protein, Piwil2. Furthermore, VPAC2 deletion leads to reduced TGFβ-1 through SP1 transcriptional regulation, linking VPAC2 to the modulation of non-tumor cell-autonomous effects in the TME. Data from *in vivo* tumor growth rates of Panc02 cells and knockdown of VPAC2 in the KPC.luc model confirm the role of VPAC2 signaling in promoting tumor growth and greater sensitivity to anti-PD1 therapy. These findings position VPAC2 at the intersection between tumor cell-intrinsic autocrine signaling to regulate growth and paracrine signaling to regulate immune cells in the TME.

## Materials and Methods

### Cell lines and reagents

MT5 and KPC.luc cells were generous gifts from Dr. Tuveson and Dr. Logsdon, respectively, and Panc02 cells were provided by Dr. Pilon-Thomas. MT5 was cultured in 1XRPMI medium and KPC.luc and Panc02 in 1XDMEM. All media were supplemented with 10% FBS, 10 mM glutamine, and antibiotics. CRISPR/Cas9 VPAC2 knockout pools for Panc02 were obtained from Synthego. KPC.luc cells were transduced with murine VPAC2 shRNA lentiviral particles (Santa Cruz Biotechnology) at a Multiplicity of Infection (MOI) of 20. CRISPR-KO VPAC2KO Panc02 cells were similarly transduced with VPAC2-overexpression lentiviral particles (Origene) at 50 MOI for the rescue experiment. Genetically modified cells were selected using puromycin and single-cell cloned via limiting dilution.

### Mice

All experimental procedures were approved by the Institutional Animal Care and Use Committee at Emory University. Female or male C57BL/6 (000664, the Jackson Laboratory) at 8-10 weeks of age were used for *in vivo* experiments. All animals were maintained according to guidelines, *The Guide for Care and Use of Laboratory Animals* (National Research Council).

### Patients and samples

Formalin-fixed paraffin-embedded (FFPE) tissue blocks from pancreatic cancer patients were obtained under an IRB-approved protocol [IRB00102595] at the Winship Cancer Institute of Emory University.

### Cell proliferation and Colony formation assay

Cell viability was assessed using the MTT cell proliferation kit I (Roche), following the manufacturer’s instructions. The percentage of cell viability was normalized to 0 hr. For colony formation assays, 50-200 cells were plated in triplicates in 6-well plates and grown over 8 to 10 days. Following culture, cells were stained with 0.5% crystal violet for 30 minutes at room temperature (RT). Images were acquired and analyzed using CountPHICS. Colonies were manually counted for experiments if the average number of visible colonies for the groups was less than 50.

### In vitro drug treatment

1mM stock of VPAC2-specific antagonist (Pg 99-465 Trifluoroacetate, Bachem) and SP1 inhibitor (Plicamycin, MedChemExpress) were prepared in pyrogen-free water and DMSO respectively. Cells were treated with 10μM VPAC2 antagonist or 0.2-1μM plicamycin daily.

### RNA Sequencing analysis of parental versus VPAC2 CRISPR-KO cells

RNA Sequencing and subsequent analyses were performed by Azenta Biosciences. Briefly, 1 million Panc02 cells were pelleted and submitted for bulk RNA-Sequencing analysis on the Illumina HiSeq 4000 platform. DESeq2 analysis on normalized gene counts was performed, followed by the Wald test to generate p-values and log2 fold changes. Genes with adjusted p-values of less than 0.01 and relative expression levels of log2 greater than -1 and 1 were considered significant. GSEA analysis was performed using Board Institute Hallmarks mouse gene sets on differentially expressed genes. Volcano plot and dot plots were generated using R-studio (Version 4.3.1).

### Reverse Transcriptase Polymerase Chain Reaction (RT-PCR) and qRT-PCR

Total RNA was isolated using RNeasy Plus Micro Kit (Qiagen), and first-strand cDNA was prepared using an AMV RNA PCR kit (TaKaRa) from 1 mg of total RNA. PCR amplification for VPAC2 mRNA detection was carried out as previously described (9). For qPCR, PowerUp™ SYBR™ Green Master Mix was used. The PCR reaction was carried out in 96-well plates using the 7500 Fast Real-Time PCR System (Applied Biosystems). A melting curve analysis was performed for each sample to verify PCR specificity, and no template samples were used as a negative control. Fold changes were calculated by the ΔΔCt method by using GAPDH as a housekeeping gene. Primers used are detailed in Supplementary Table 2.

### TGFβ multiplex immunoassay

0.3 × 10^6^ KPC.luc, Panc02, and MT5 were cultured in 6 well plates with 2 ml of respective media. Cell-free media collected 72 hours after culture were tested for 3 TGF beta isoforms using the U-plex TGFβ Combo cytokine plate for mouse (MSD, Cat. K15242K-1). Experiments were performed by Emory Multiplexed Immunoassay Core (Emory University).

### T cell activation and function

Spleens were harvested from naïve C57BL/6 mice. T cells from splenocytes were isolated using a mouse pan-T cell isolation kit (Cat. 130-095-130, Miltenyl Biotec). 100,000 to 200,000 T cells were seeded in 96-well flat bottom plates coated with 0.1 μg/ml anti-mouse CD3e antibody (Cat. 16-0031-86, Invitrogen™) and cultured in PDAC conditioned media containing 30U/ml recombinant mouse IL-2 for 24 to 72 hours . For cytokine analysis, Golgi plug (Cat. 51-2301KZ, BD) and Golgi stop (Cat. 51-2092KZ, BD) were added 4 hours before collection for flow cytometry analysis.

### Transcription Factor Profiling

Nuclear proteins were isolated from 70-80% confluent KPC.luc cells using Nuclear Extraction Kit (Cat. NPBP-29447, Novus) and quantified using Bradford protein assay. 9μg of total nuclear proteins from KPC cells was subjected for active transcription factor profiling (Cat. FA-1002, Signosis Inc.). Data was normalized to each plate using luminescence values from Gli-1 transcription factor that did not change between groups.

### In vivo efficacy studies

For the subcutaneous model, 1 × 10^6^ KPC.luc or Panc02 were injected near the right flank of female or male C57BL/6 mice. The study reached endpoint when the tumor volume reached 500mm^3^ or if the tumor was ulcerated. For NOD.Cg-Prkdc^scid^ Il2rg^tm1Wjl/^SzJ (NSG) immunocompromised mice, 1 × 10^6^ Panc02 or KPC.luc were injected subcutaneously and followed until tumor volume reached 1000mm^3^ and/or ulcerated. Vernier calipers were used to measure the tumor dimensions and the tumor volume was calculated using the formula: tumor volume = ½*(length × width × height). For the orthotopic KPC.luc model, mice were anesthetized, and the 200,000 KPC.luc cells were suspended in PBS: Matrigel (1:1) and injected in the tail of the pancreas following laparotomy. Tumor growth was monitored weekly for 24 days using IVIS bioluminescent imaging.

### In vivo treatment

C57BL/6 mice were treated with 200 μg of murine antibodies to PD-1 (Clone RMP1-14) or isotype control (Clone 2A3) for 4 days (+7, +10, +14, +17) following tumor implantation. For depletion studies, anti-CD4 (Clone GK1.5) or anti-CD8 (Clone 2.43) antibodies were injected intraperitoneally at 200 μg per mouse on days -1 and at 100 μg +1, +3, +7, +11, +15) with respect to the date of tumor implantation. Antibodies were obtained from BioXcell and are detailed in Supplementary Table 1.

### Histology

Pancreas harvested from mice with orthotopically implanted KPC.luc tumors were formalin-fixed and paraffin-embedded before being stained with H&E. All slides were dehydrated, cover-slipped, and scanned on Hamamatsu Nanozoomer 2.0 HT at 40x.

### Immunofluorescence (IF)

FFPE PDAC tissues were deparaffinized, hydrated, and underwent antigen retrieval by boiling with 1X Trilogy for 15 minutes (Cell Marque-Trilogy Buffer). Permeabilization was performed using 0.3% Triton-X-100, followed by a blocking step with eBioscience™ high protein blocking buffer for 1 hour at RT. Primary antibodies for VIP, VPAC2, and CK19 (**Suppl. Table 1**) were applied with overnight incubation at 4°C. Secondary antibodies for anti-mouse IgG (H+L) conjugated with Alexa(R) 647 and anti-rabbit IgG (H+L) conjugated with Alexa(R) 488, and anti-goat conjugated with TRITC were applied and incubated for 1 hour at RT. Tissue slides were stained with 2 μg/ml Hoechst and imaged at Plan Fluor 40X objective on a BZ-X810 epifluorescence microscope (Keyence Corp). Multiple images were acquired and stitched using Keyence’s image stitcher function and merged into a composite image using Fiji.

### Statistical Analysis

For the comparison of one variable in more than two groups, one-way ANOVA followed by Dunnett’s multiple comparison post-hoc test or Welch’s correction was used. For two groups comparison, an unpaired T test with Welch’s correction or Mann-Whitney test was used. Two-way ANOVA followed by Bonferroni’s multiple comparisons test was used for multi-variable group analysis. For survival data, the Kaplan-Meier method with log-rank tests was used and the hazard ratio was estimated by Cox proportional hazard model. R-squared values were generated from a linear regression model for PDAC TCGA dataset. P values less than 0.05 were considered significant. All statistical analyses were conducted using GraphPad Prism software, version 10.1.0.

## Data Availability Statement

Bulk RNA Sequencing data is available under the GEO accession number (pending). The data generated in this study are available upon request from the corresponding author.

## Results

### VIP and VPAC2 are co-expressed in human and murine pancreatic ductal adenocarcinoma

To study the role of the VPAC2 receptor in PDAC, we first examined VPAC2 expression in human PDAC tissue and found VPAC2 colocalization with the cancer epithelial cell marker, cytokeratin 19 (CK19) (**Fig.1A**). Moreover, we found that CK19 positive PDAC cells co-expressed VPAC2 with VIP (**Fig.1B, Suppl. Fig.S1A**). The relationship between VIP and VPAC2 was corroborated using a PDAC dataset from the Cancer Genome Atlas (TCGA) (n=149), where VIP mRNA expression was positively correlated with VPAC2 mRNA expression (R^2^=0.34, p<0.0001) but not correlated with VPAC1 mRNA expression (**Fig.1C and D**). We found that VPAC2 mRNA expression trended towards higher median expression in Stage III/IV patients compared to Stage I and II. However, few stage III/IV patients are reported in the dataset (**Suppl. Fig.S1B**). Across the various point mutations on K-ras, that occur frequently in tumors from PDAC patients, we observed a similar distribution of expression of VPAC2 mRNA in K-ras-mutant versus wild type K-ras, but a trend of lower median expression in the mutant K-ras tumors (**Suppl. Fig.S1C**). VPAC2 levels did not differ significantly between males and females (**Fig.1E**). To further study the clinical prognostic value of VIP and VPAC2, patients were stratified as Low_VIP versus High_VIP (cut-off value at 56, expression range 0 to 2149) and Low_VIPR2 versus High_VIPR2 (cut-off value at 22, expression range 0 to 297). The cut-off values for high and low expression were determined using methods previously published (22). Although multiple factors can impact patients’ relapse-free survival (RFS) (23,24), we found significantly higher RFS for patients with low expression of both VIP and VPAC2 (Low_VIP_VIPR2; 50 months) than patients with high VIP and VPAC2 (High_VIP_VIPR2) (18 months; HR:1.92, P <0.05), regardless of disease stage (**Fig.1F, Supp. Fig.S1D)**. On the other hand, there was no significant difference in RFS for patients with low VIP and VIPR1 versus high VIP and VIPR1 (**Supp. Fig.S1E**). These data suggested the hypothesis that autocrine signaling of VIP via the VPAC2 receptor may regulate PDAC tumorgenicity.

**Figure1.**
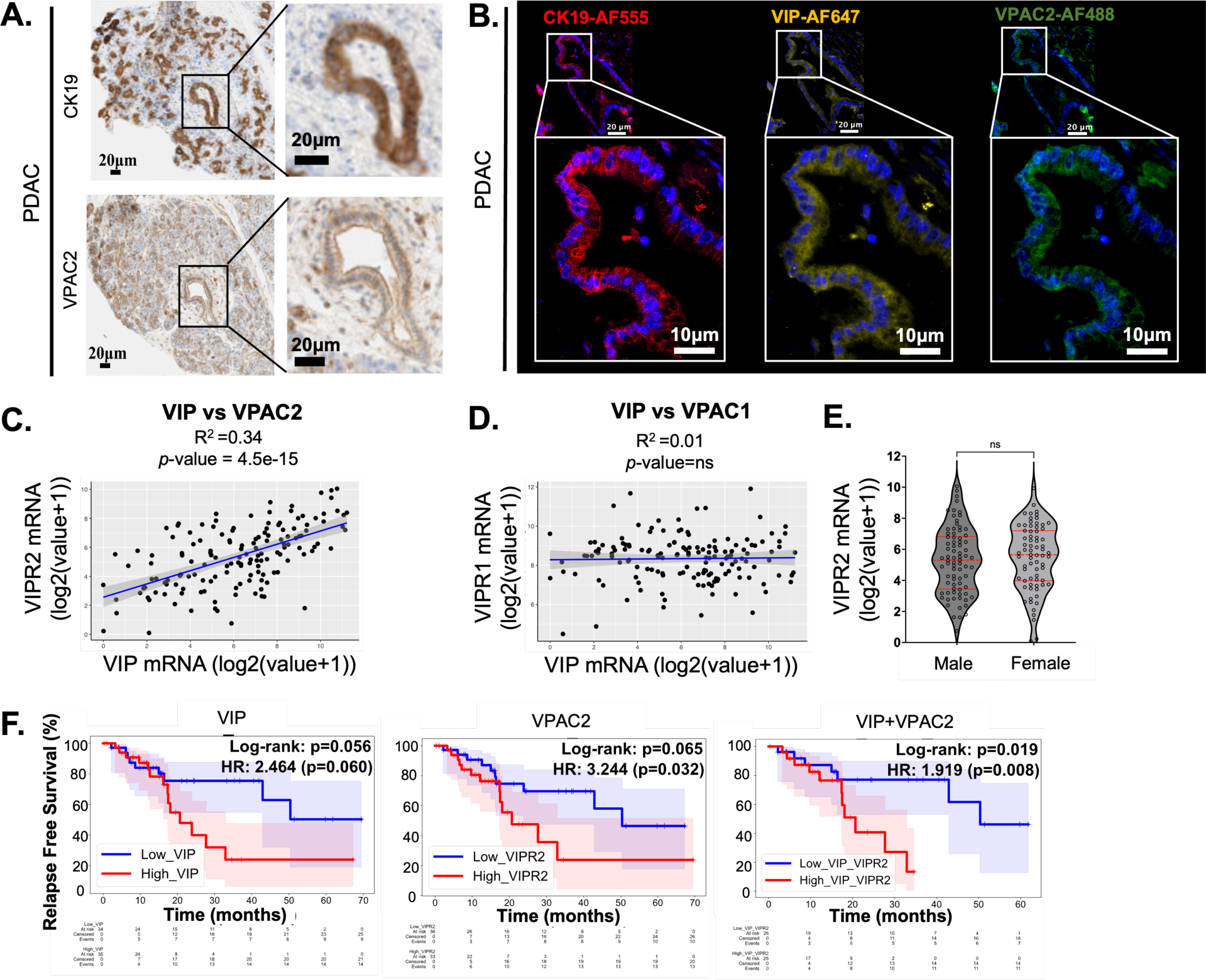
VIP and VPAC2 is co-expressed in pancreatic ductal adenocarcinoma. **(A)** Immunohistochemistry staining on PDAC tissue for VPAC2 and cytokeratin 19 (CK19). (**B**) Immunofluorescence on PDAC tissue showing expression of VIP (Mouse anti-VIP antibody), VPAC2 (Rabbit anti-VPAC2) and CK19 (Goat anti-CK19). Anti-Goat Alexa (R) 555, anti-mouse Alexa(R) 647 and anti-rabbit Alexa (R) 488 secondary antibodies was used for fluorescence detection. CK19 staining shown for pancreatic cancer ductal cells. (**C**) Linear regression model for *VIP* and *VIPR2* (encoding VPAC2) and between **(D)** VIP and *VIPR1* (encoding VPAC1) from TCGA Pancreatic Cancer dataset (n=149). **(E)** VPAC2 mRNA expression between male and female PDAC patients, extracted from TCGA Pancreatic Cancer dataset. **(F)** Relapse free survival for PDAC patients with high and low expression of VIP and VPAC2. The shaded colors below and above the survival curves correspond to 95% confidence interval for the respective patient groups. For survival analysis, Log rank test was performed for survival difference between patients. Hazard ratio (HR) was estimated by Cox proportional hazard model.

### Decreased colony formation from disruption of VPAC2 on cancer cells is VIP dependent

To explore the cell-intrinsic role of VPAC2 signaling on PDAC growth, we performed CRISPR-Cas9 deletion of VPAC2 on a murine PDAC cell line, Panc02. Lack of VPAC2 mRNA and VPAC2 protein in Panc02 cells, from now on termed VPAC2 knock out (VPAC2KO) (**Fig.2A, Suppl. Fig.S2A and B**), led to a 1.9-fold reduction in downstream CREB phosphorylation (47.4%±7.5 vs. 25.0%±5.5 in parental vs. VPAC2KO respectively) (**Fig.2B**) indicating the suppression of VIP/VIP receptor pathway, absent in CRISPR-control (**Suppl. Fig.S2C**). The VPAC2KO cells had a slightly delayed *in vitro* proliferation compared to parental cells, while no difference was observed between parental and CRISPR-control cells (**Fig.2C, Suppl. Fig.S2D**). Delayed proliferation was observed at 24 hours, but growth rates were similar at later time points. We hypothesized that VPAC2KO cells may have impaired growth at low confluency. To test this hypothesis, we plated the cells at 100 cells per 6-well plate (9.6 cm^2^/well) and found significantly fewer colonies with reduced size in VPAC2KO versus parental cells (27±4 vs. 62±7 colonies, respectively; **Fig.2D**), which were rescued upon VPAC2-overexpression (VPAC2-ORF) in VPAC2KO cells (37±5 colonies in control-ORF vector and 69±8 in rescue VPAC2-ORF; **Fig.2E and F, Suppl. Fig.S2E**). Reduced colonies were also observed by blocking VPAC2 pharmacologically with a VPAC2-specific antagonist in cultured Panc02, and MT5 murine PDAC cell lines, as well as following transient knockdown of VPAC2 using siRNA (**Fig.2H and I, Suppl. Fig.S2F and G**). Furthermore, neutralizing VIP in the supernatant using an anti-VIP antibody led to significantly reduced colonies, similar to VPAC2 antagonist treatment (**Fig.2H**).

**Figure2.**
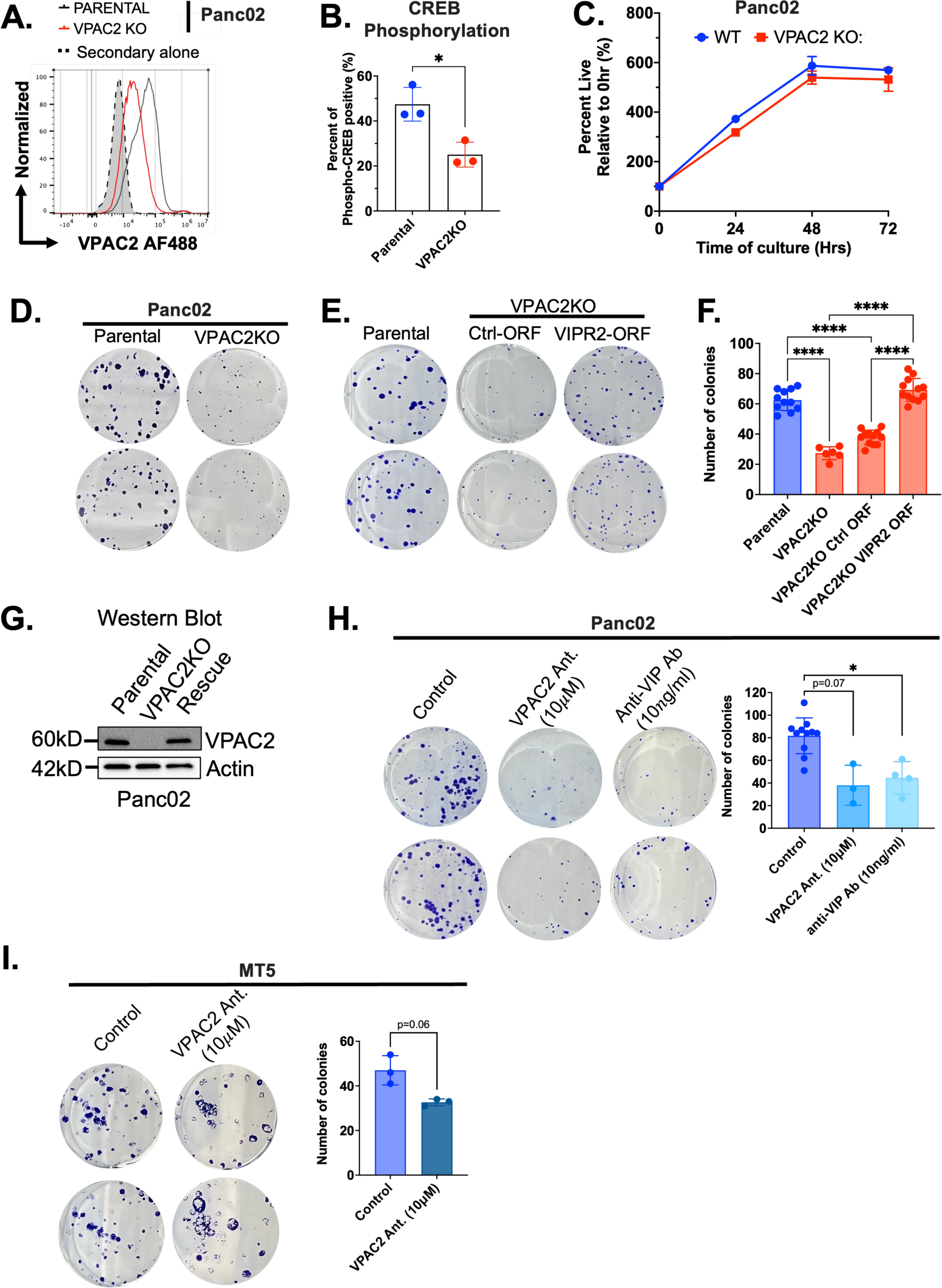
Absence of VIP and VPAC2 signaling leads to decreased colony formation in vitro. **(A)** Flow cytometry for surface expression detection of VPAC2 using rabbit polyclonal anti-VPAC2 antibody and **(B)** levels of phosphorylated CREB in parental Panc02 versus CRISPR-edited VPAC2KO Panc02 cells. **(C)** MTT assay showing growth of Panc02 cells over 72 hours. **(D)** Representative pictures of crystal violet colony assay for parental Panc02 and VPAC2KO Panc02 cells and **(E)** VPAC2-ORF rescue Panc02 cells. Cells were stained with crystal violet stain after 8 days of culture. **(F)** Number of colonies as computed using countPHICS software. **(G)** Western blot confirmation of VPAC2KO and VPAC2 rescue in VPAC2KO Panc02 cells using polyclonal anti-VPAC2 antibody. Crystal violet colony assay for **(H)** Panc02 and **(I)** MT5 treated with VPAC2-specific peptide-based antagonist (10μM) and anti-VIP antibody (10ng/ml). Cells were treated daily for 8 days with VPAC2 antagonist whereas treated only once with anti-VIP antibody. Data are presented as bar graphs or line plots ± standard deviation (SD). For B and I, two-tailed unpaired t test was used. For F and H, one-way ANOVA test following by Dunnet’s multiple comparison post-hoc test was performed. *p<0.05, ****p<0.0001.

### Decreased expression of *Piwil2*, a stem-cell-related gene downstream of VPAC2 signaling, decreased *in-vitro* colony formation

To further examine the downstream consequences of VPAC2 signaling on pancreatic cancer cells, we performed bulk RNA sequencing on parental, CRISPR-control, and VPAC2KO Panc02 cells. Principal Component Analysis (PCA) indicated separate clustering for the parental vs. VPAC2KO samples, but not for parental vs. CRISPR-control, which highlights the differences in a broad range of expressed genes between parental vs. VPAC2KO cells (**Suppl. Fig.S3A and B**). To account for genes in which expression may have been non-specifically affected due to CRISPR-Cas9 editing, we removed genes differentially expressed between parental and CRISPR-control samples (**Suppl. Fig.S3C**) from the differential gene analysis between parental and VPAC2KO. We found a total of 371 genes downregulated and 301 genes upregulated genes in the VPAC2KO compared to parental cells (**Fig.3A**). Among the downregulated genes, Piwi Like RNA-Mediated Gene Silencing 2, commonly known as *Piwil2*, was significantly downregulated compared to parental cells (log2 fold change -5.6; **Fig.3A**). We confirmed mRNA levels using qPCR, demonstrating reduced Piwil2 mRNA in the VPAC2KO cells, which increased by 2.8-fold following rescue of VPAC2 expression in VPAC2KO Panc02 cell line (**Fig.3B**). We also observed a decrease in Piwil2 mRNA in the KPC.luc and MT5 cell line after blocking VPAC2 signaling with a VPAC2 antagonist (**Fig.3C**). To test the role of Piwil2 in survival of PDAC cells, we performed siRNA transfection targeting Piwil2. We found decreased colony formation compared to control transfected cells (**Fig.3D, Suppl. Fig.S3D**). Importantly, VPAC2 mRNA expression was positively correlated with Piwil2 mRNA expression in the human PDAC dataset (R^2^ = 0.1, p<0.01; **Fig.3E**). Additionally, we identified other gene sets that were significantly downregulated in the VPAC2KO clone using Hallmarks gene sets GSEA analysis and observed myc targets as one of the significantly downregulated pathways compared to parental cells (**Fig.3F, bold**).

**Figure 3.**
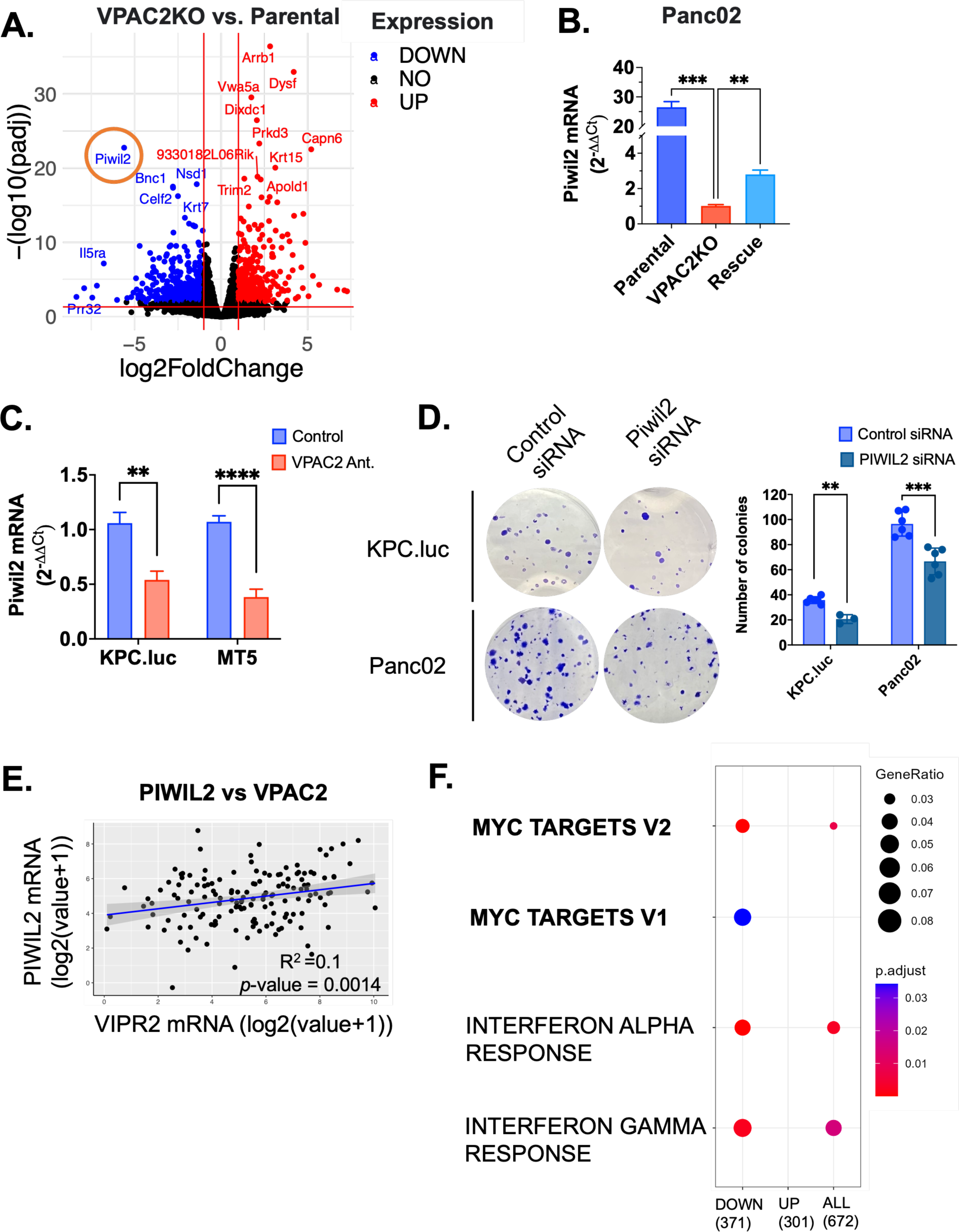
Decreased stem-cell related gene, *Piwil2*, downstream of VPAC2 signaling leads to decreased colony formation *in-vitro*. **(A)** Volcano plot of significantly downregulated (blue) and upregulated (red) genes between parental and VPAC2KO Panc02 cells from RNA Sequencing analysis. Piwil2 (circled in orange) were one of the top 50 genes downregulated in the VPAC2KO compared to parental cells. **(B)** qRT-PCR confirmation of Piwil2 mRNA expression in VPAC2KO and VPAC2-rescue Panc02 cells versus parental Panc02 cells (n=5). **(C)** qRT-PCR for Piwil2 mRNA expression in KPC.luc and MT5 after treatment of cells for 3 days with VPAC2-antagonist *in vitro* (n=8). **(D)** Crystal violet colony assay for KPC.luc and Panc02 cells. Cells were transfected with control or Piwil2-targeting siRNA for 2 days. siRNA transfected cells were plated at 50-100 cells per 6-well plate and cultured for 8-10 days. Cells were stained with crystal violet stain after 8-10 days of culture and number of colonies were computed using countPHICS software. **(E)** Linear regression model for *VIPR2* (encoding VPAC2) and Piwil2 from TCGA Pancreatic Cancer dataset (n=149). **(F)** Hallmarks GSEA analysis on differentially expressed genes (UP and DOWN) between parental and VPAC2KO Panc02 cells. For D, data are presented as bar graphs ± standard deviation (SD). For B and C, data are presented as + standard error (SEM). For B, one-way ANOVA test following by Dunnet’s multiple comparison post-hoc test was performed. For C and D, two-tailed unpaired t test was used. *p<0.05, **p<0.01, ***p<0.001, ****p<0.0001.

### VPAC2 regulates TGFβ-1 expression via SP1 and decreases T cell function

In addition to the mRNA regulation of Piwil2 by VPAC2 receptor modulating tumor cell-intrinsic pathways, we were interested in whether the absence of VPAC2 signaling also alters the expression of cytokines that have immunosuppressive effects in the TME. Culture supernatants had low levels of the inflammation-related immunosuppressive cytokines IL6, IL10, and IL-1β, reported to be secreted by the PDAC cells (25,26), with levels of 0.5pg/ml for IL10, 3-6 pg/ml IL6, and below detection level for IL-1β in the conditioned media from three PDAC cell lines (**Suppl. Fig.S4A**). In contrast, the levels of transforming growth factor beta (TGFβ) in culture supernatants, predominantly TGFβ-1, were elevated to >1000 pg/ml (**Supp. Fig.S4B**). Given the high cytokine levels that may have an immunosuppressive role in the TME, we investigated whether VIP/VPAC2 signaling alters TGFβ levels. We first confirmed that deletion of VPAC2 does not impact the expression of VIP as a potential positive feedback loop (**Suppl. Fig.S4C and D**). Knock out of VPAC2 in Panc02 (VPAC2KO) or knockdown of VPAC2 (VPAC2KD) in KPC.luc led to decreased expression of TGFβ-1, but not TGFβ-2 and TGFβ-3 (**Fig.4A, Suppl. Fig.S4E and F**). Knock-down validations for VPAC2 in KPC.luc cell line are shown in supplementary Figures 3G & H. In addition, treatment with the VPAC2 antagonist also led to decreased TGFβ-1 levels (**Fig.4B**). The decrease of TGFβ-1 in VPAC2KO Panc02 cells did not have consistent effects on the EMT-related proteins (data not shown), which led us to test if the secreted TGFβ-1 had paracrine effects on T cells. We cultured naïve T cells in PDAC-conditioned media from parental, VPAC2KO, and VPAC2-ORF rescue and assessed the effect of secreted factors on T cells (**Fig.4C**). T cells cultured in VPAC2KO-conditioned media (VPAC2KO-CM) had higher activation levels compared to parental-conditioned media (parental-CM) as measured by CD69 expression at 24 hours (22.8%±1.7 vs 18%±0.7) (**Suppl. Fig.S4G**). Moreover, the CD8+ T cells cultured in VPAC2KO-CM had greater IFNγ (3.4% ± 0.3 vs 0.2% ± 0.2) and TNFα expression (9.3%±1.9 vs. 1.7%±0.4) than parental-CM (**Fig.4D**). We also saw greater T cell proliferation following 6 days of culture in VPAC2KO-CM than parental-CM (33.5% ± 4.1 vs. 5.0% ± 0.7 CFSE^low^; **Suppl. Fig.S4H**). The increased CD69, IFNγ, and TNFα expression on T cells cultured in VPAC2KO-CM were abrogated or reduced to levels similar to parental-CM when cultured in VPAC2-ORF rescue-conditioned media (Rescue-CM) (**Fig.4D, Suppl. Fig.S4G-J**). Addition of recombinant TGFβ-1 to VPAC2KO-CM also led to similar T cell phenotypes as parental-CM, thereby linking TGFβ-1 mediated T cell suppression to VPAC2 signaling in PDAC cells (**Fig.4D, Suppl. Fig.S4I and J**). To elucidate the regulation of transcription factors downstream of VPAC2 signaling, we performed transcription factor profiling using nuclear proteins extracted from control and VPAC2 knock-down KPC.luc cells. We found reduced levels of GR/PR, SP1, ELK, OCT4, p53 and Brn-3 in the absence of VPAC2 (**Fig.4E**). We focused on the SP1 transcription factor given previous work indicating SP1 binding to the promoter region of TGFβ-1 for its regulation (27–30). We validated the reduced nuclear expression of SP1 via western blot in both KPC.luc VPAC2KD and Panc02 VPAC2KO clones compared to control cell lines (**Fig.4F**). Furthermore, we found that treatment of PDAC cells with SP1 inhibitor, plicamycin, decreased TGFβ-1 (**Fig.4G**), representing a novel regulation of TGFβ-1 by SP1 downstream of VPAC2 signaling.

**Figure 4.**
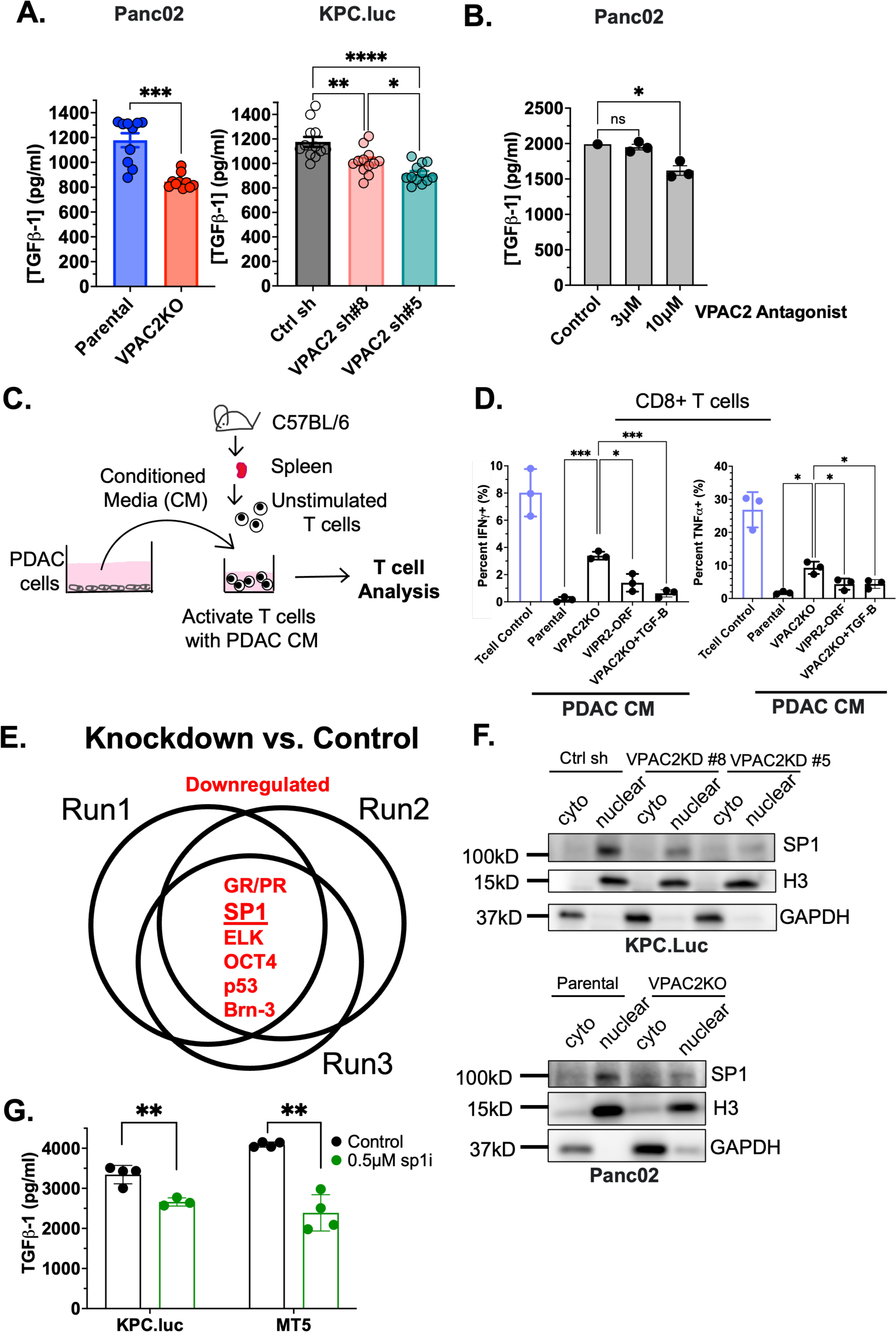
Disruption of VPAC2 pathway leads to reduced TGFβ-1 via SP-1 and increases T cell function. **(A**) Levels of secreted TGFβ-1 in Panc02 and KPC.luc cultures. CRISPR-KO or lentiviral knock down of VPAC2 cells were compared to control cultures. **(B)** TGFβ-1 secretion in Panc02 upon treatment with 3μM and 10μM of VPAC2-antagonist for 3 days. **(C)** Schematic diagram for T cell assay performed with conditioned media from PDAC-cells. **(D)** Expression of IFNγ and TNFα on CD8+ T cells following culture in conditioned media from parental, VPAC2KO, VPAC2-rescue Panc02 cells. Complete RPMI media was used as T cell control media. Recombinant TGFβ-1 at 1ng/ml was added to conditioned media from VPAC2KO cells as additional group. T cells were stimulated with 0.1μg/ml anti-CD3 coated plates and cultured for 72 hours. Golgi plug and Golgi stop was added to culture 4 hours prior to staining for IFNγ and TNFα cytokine by flow cytometry (n=3). **(E)** Venn diagram for commonly downregulated transcription factors between control and VPAC2KD #8 KPC.luc cells (n=3). The downregulated transcription factors shown in red are ranked in order based on highest to lowest fold change, SP1 ranking second (underlined). **(F)** Western blot validation of reduced SP1 expression in the nucleus of two VPAC2KD clones (#8 and #5) in KPC.Luc cells and VPAC2KO Panc02 compared control transduced and parental cells respectively. Histone 3 (H3) protein shown as loading control for nuclear extract and GAPDH for cytoplasmic extract. **(G)** Levels of secreted TGFβ-1 in KPC.luc and MT5 cell line following treatment with sp1 inhibitor, plicamycin, at 0.5μM for 72hours. For A, data are presented as bar graphs ± standard error (SEM). For B,D, and G, data are presented as ± standard deviation (SD). For A, D and G, unpaired t test with Welch’s correction was used. For B, one-way ANOVA test following by Dunnet’s multiple comparison post-hoc test was performed. *p<0.05, **p<0.01, ***p<0.001, ****p<0.0001.

### The absence of VPAC2 decreases tumor growth and increases sensitivity to aPD-1 therapy in the subcutaneous PDAC model

Next, we tested if the *in vitro* findings of tumor-intrinsic and tumor-extrinsic effects of VPAC2 signaling were recapitulated *in vivo*. We first inoculated control or VPAC2KD KPC.luc to C57BL/6 mice and observed significantly slower growth rates in mice with VPAC2KD tumors (**solid lines, Fig.5A**). The control transduced KPC.luc cell line (control sh) grew progressively in C57BL/6 mice (**Fig.5A, black line**). The two clones knocked down for VPAC2 (VPAC2 sh #5 and #8) grew at comparable rates to control sh tumors until day 10. At that time, the tumors began to shrink, leading to a significantly higher fraction of tumor-free mice at the end of the study (2/9 for mice with control sh tumors versus 6/10 and 7/10 mice with VPAC2 sh tumors) (**Fig.5B and C**). Tumor regression in the absence of VPAC2 was not observed in the NSG model, where there was only slight delay in growth, suggestive of modest cell-intrinsic growth effects (**dashed lines, Fig.5A and B**).

**Figure 5.**
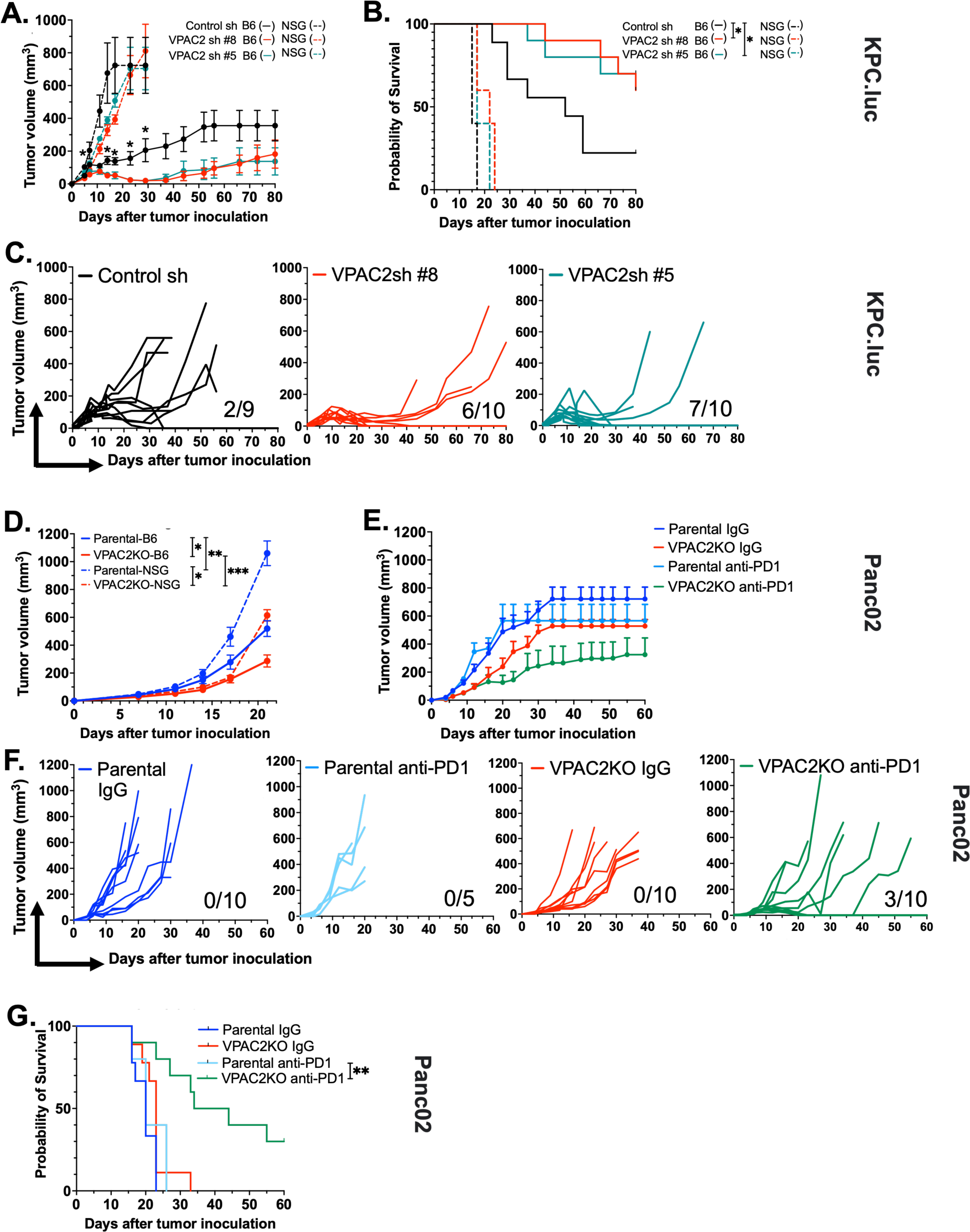
Absence of VPAC2 leads to decreased tumor growth in vivo in a tumor intrinsic and extrinsic manner in subcutaneous PDAC models. **(A)** Average tumor volume over time for control sh and VPAC2sh KPC.luc subcutaneously injected to C57BL/6 (n=9-10) (solid line) and to NOD SCID gamma (NSG) mice (n=5) (dashed line). **(B)** Kaplan-Meir survival plots for mice corresponding to A. **(C)** Spider plots showing tumor volume for individual C57BL/6 mouse injected with control (control sh, n=9) and VPAC2 knock down (VPAC2sh, n=10 each clone) KPC.luc cells (Top Panel). Numbers indicate the fraction of C57BL/6 mice that were tumor free at day 80 post tumor implantation. **(D)** Average tumor volume over time for parental and VPAC2KO Panc02 cells subcutaneously injected to C57BL/6 (n=15-20) (solid line) and to NOD SCID gamma (NSG) mice (n=5) (dashed line). **(E)** Average tumor volume over time for parental and VPAC2KO Panc02 cells injected to male C57BL/6 (n=5-10) and treated with isotype IgG or anti-PD1 antibody. **(F)** Spider plots showing tumor volume for individual male C57BL/6 mouse from E. Numbers indicate the fraction of C57BL/6 mice that were tumor free at day 60 post tumor implantation. **(G)** Kaplan-Meir survival plots corresponding to E. Tumor volumes were measured by vernier calipers one to two times a week until study endpoint. Data are presented ± standard error (SEM). For A and D, two-way ANOVA with Bonferroni correction was used. Log-rank test was used for statistical differences for Kaplan-Meier curves. *p<0.05, **p<0.01, ***p<0.001.

**Figure 6.**
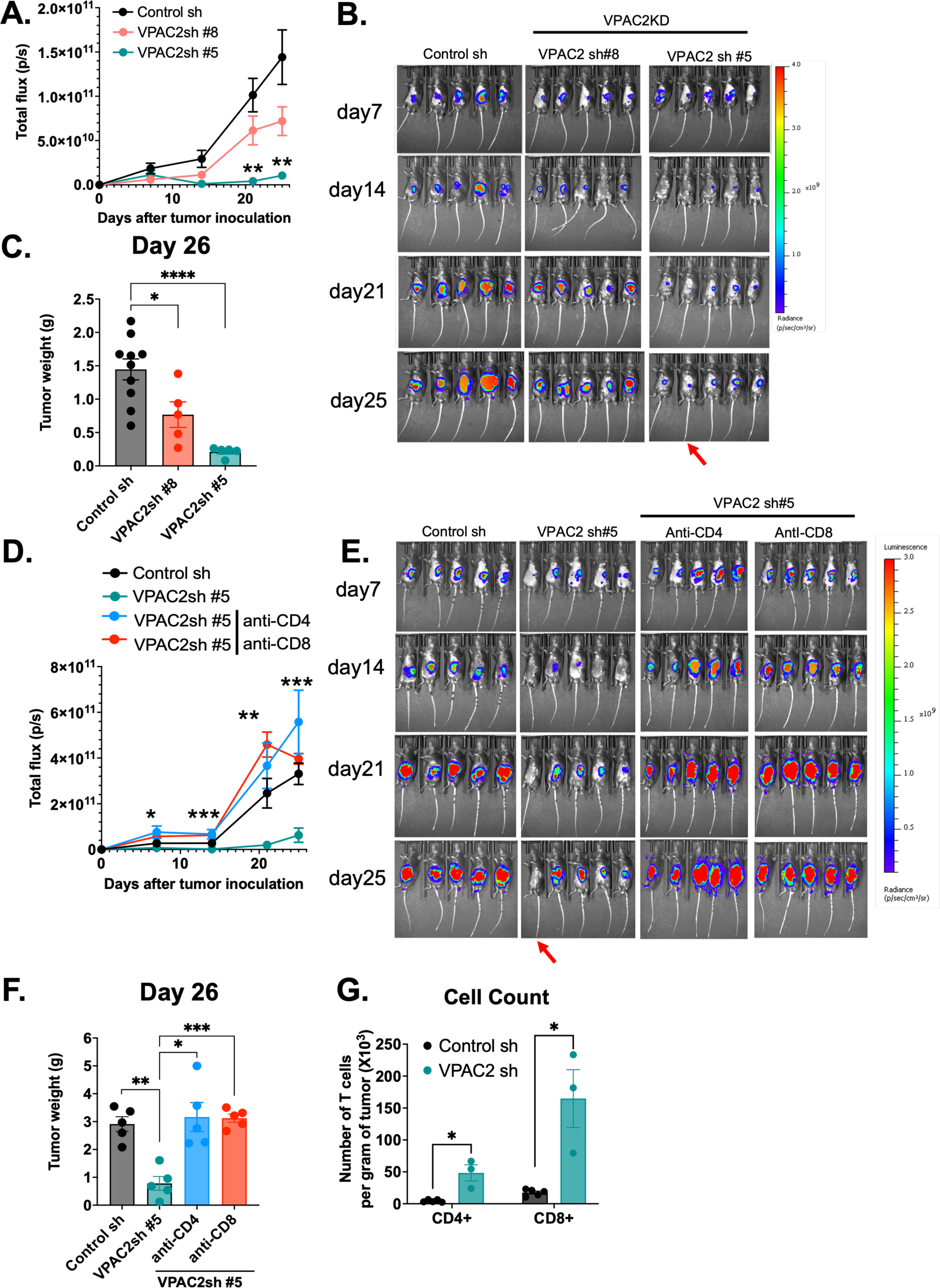
Knock down of VPAC2 on cancer cell reduces T-cell dependent tumor burden in KPC.luc orthotopic model. Control sh and VPAC2 sh (#8 and #5) KPC.luc cells were surgically implanted in the tail of the pancreas of C57BL/6 mice. **(A)** Total flux monitored over time by IVIS bioluminescent imaging. Isoflurane was used for anesthesia for bioluminescent imaging. **(B)** Pictures from IVIS bioluminescent imaging acquired weekly for the individual mice implanted with control or VPAC2KD KPC.luc cells for 25 days. Red Arrow indicating mice with minimal tumor residual at study end point. **(C)** Bar graph showing weight of tumor in grams on day 26 when the mice were euthanized. **(D)** C57BL/6 mice implanted with VPAC2sh #5 clone KPC.luc receiving either monoclonal CD4 or CD8 monoclonal antibody compared to isotype treatment. Total flux from IVIS imaging shown comparing the groups. **(E)** Pictures from IVIS bioluminescent imaging acquired weekly for the individual mice implanted with control or VPAC2KD KPC.luc with isotype or anti-CD4/anti-CD8 blockade. Red Arrow indicating mice with minimal tumor residual at study end point. **(F)** Bar graph showing weight of tumor in grams on day 26 when the mice were euthanized corresponding to groups in D-E. Upon euthanasia, KPC.luc tumors were dissociated as single cells and analyzed for T cell count by flow cytometry. **(G)** Bar plot showing percent of T cell count as gated on CD4+ or CD8+. All data are presented as ± standard error (SEM). For A, and D, two-way ANOVA with Bonferroni correction was used. For C and F, one-way ANOVA test following by Dunnet’s multiple comparison post-hoc test was used. For G, Mann-Whitney test was performed. *p<0.05, **p<0.01 and ***p<0.001, ****p<0.0001.

Similar delayed growth was observed in the VPAC2KO Panc02 model. However, all mice succumbed to the tumor with faster tumor growth rates in NSG mice than C57BL/6 mice (**Fig.5D, Suppl. Fig. S5 A**). Notably, tumor volume at day 21 was higher in parental vs VPAC2KO Panc02 tumors in NSG mice, but parental tumors did not have higher expression of Ki67, a proliferation marker (**Suppl. Fig.S5B-D**). With *in vitro* experiments showing that VPAC2 expression on PDAC cells alters T cell phenotype via tumor-cell-extrinsic mechanisms, we investigated the response of parental vs VPAC2KO Panc02 tumors to anti-PD-1 therapy. On average, the tumor growth rate of the parental cells was not altered by anti-PD1 therapy, consistent with previous reports (31,32), whereas anti-PD1 therapy suppressed tumor growth in mice with VPAC2KO tumors (**Suppl. Fig.S5E**). The median survival of mice challenged with the parental tumor was 27.5 days versus 50.5 days in those with VPAC2KO tumors following anti-PD1 therapy, albeit not significant (*p*=0.2, **Suppl. Fig.S5F**). The survival advantage was significant in male mice, that have less anti-cancer immunity than female mice (33), in which no mice with parental tumors were rendered tumor-free with anti-PD1 therapy (0/5) versus 30% tumor-free mice (3/10) with VPAC2KO tumors (**Fig.5E and F**). No significant differences in growth rates or survival were seen comparing parental versus VPAC2KO cells in female C57BL/6 mice (**Suppl. Fig.S5G and H**).

### The absence of VPAC2 reduces tumor burden in a clinically relevant orthotopic model and increases T cell number and function in the tumor microenvironment

The significant effects of VPAC2 in controlling the growth of subcutaneous KPC.luc in C57BL/6 mice led us to further investigate the role of VPAC2 signaling in a more clinically relevant orthotopic PDAC model. We surgically implanted control sh or VPAC2 sh clones of KPC.luc in the tail of the pancreas and monitored tumor growth *in vivo* by bioluminescence (BLI). We documented successful tumor engraftment 7 days following surgery (**Suppl. Fig.S6A**). In contrast to the control sh KPC.luc tumors, the VPAC2 knocked down clones (VPAC2 sh #8 and #5), show reduced tumor growth rates (**Fig.5A**). In fact, we found that one of the mice with initial engraftment of the VPAC2 sh#5 tumor had no detectable residual tumor at necropsy (**Fig.5B, red arrow**). These findings were further supported by lower tumor weights at day 26 for VPAC2 sh clones versus control sh KPC.luc tumors (tumor weights for control sh > VPAC2 sh#8 > VPAC2 sh#5) with no evident histological differences in the tumors after H&E staining (**Fig.5C, Suppl. Fig.S6B**). We did not observe any lung or liver metastasis at day 26 post tumor implantation in this model (**Suppl. Fig.S6C**). To test whether the increased tumor control in the VPAC2 knock down KPC.luc model was dependent on CD4+ or CD8+ T cells, we next performed depletion studies with anti-CD4 and anti-CD8 antibodies (**Suppl. Fig.S6D**). Once again, one of the mice transplanted with the VPAC2KD clone in the isotype treatment group did not have a progressive tumor and had histologically normal pancreas tissue at necropsy (**Fig.5E, red arrow**). On average, mice with the VPAC2KD tumors treated with isotype IgG had smaller tumors than control tumors. In contrast, treatment with anti-CD4 or anti-CD8 blockade led to the loss of tumor control as measured by increased total BLI flux and tumor weights (**Fig.5D-F**). Confirmation that the antibody treatment depleted CD4+ and CD8+ in the spleen and tumor at the end of the study are shown in Supplementary Figure 6E. Finally, we observed an increased number of CD4+ and CD8+ T cells infiltrating into the VPAC2 knock-down KPC.luc tumors compared to control tumors (**Fig.5G**) with significantly fewer PD1^high^ CD8+ T cells but more Ki67 positive CD8+ at the tumor site (**Suppl. Fig.S6F-I**).

## Discussion

PDAC remains a challenging cancer with limited treatment success, due in part to its desmoplastic and immunosuppressive TME, constituting of a rich stromal compartment comprised of extracellular matrix, cancer-associated fibroblasts, immunosuppressive cell types such as MDSCs, regulatory T cells, and regulatory B cells, among others (1,6,7,34). In this study, we identify the VPAC2 signaling pathway as a novel vulnerability of PDAC involved in tumor growth and the generation of an immunosuppressive TME.

In tumors from PDAC patients and murine models of PDAC, we have previously demonstrated overexpression of VIP (9). Although, there have been other reports on VIP-receptor (VIP-R) signaling in PDAC, much of the prior work involved receptor activation with VIP or weak antagonism by VIPhyb (19–21). However, the downstream signaling pathways of VIP signaling on PDAC cells via VPAC1 and/or VPAC2 receptors remain to be elucidated. While studies have reported the expression of both VPAC1 and VPAC2 on human tumors (35,36), we observed a strong positive correlation of VIP expression with VPAC2, not VPAC1 expression in the PDAC TCGA dataset, leading us to hypothesize that VIP signals through VPAC2 on PDAC cells. We demonstrated that VPAC2 regulates tumor-intrinsic signaling by increasing tumor growth *in vitro* and *in vivo*. Mechanistically, VPAC2 increases stem-cell-related Piwil2, the blockade of which leads to reduced PDAC cell proliferation *in vitro.* While several studies have demonstrated the tumor-promoting role of Piwil2 (37,38), there is no direct report of Piwil2 on the growth of PDAC cells. PIWI proteins interact with non-coding RNAs, commonly known as (piRNAs), in the nucleus, to regulate stem-cell maintenance and self-renewal genes (39). Amongst various genes, Piwil2 promotes c-myc expression by facilitating NME/NM23 nucleoside diphosphate kinase 2 to bind to the c-myc promoter and subsequently enhance tumor cell proliferation (40). In accordance with the previous publication (40), GSEA analysis showed myc targets significantly downregulated in VPAC2KO cells compared to parental cells, which correlates with the downregulation of Piwil2. Alternatively, Piwil2 can use epigenetic gene regulation to suppress apoptosis and promote cancer growth (41). Further mechanistic studies on how Piwil2 specifically mediates tumor-cell intrinsic growth following VPAC2 signaling in PDAC are necessary. However, these data together suggest the importance of the VIP/VPAC2 axis on positive regulation of tumor cell growth via Piwil2, thus implicating the autocrine signaling of VPAC2 on PDAC cells in a cell-intrinsic manner.

Beyond its tumor intrinsic role, VPAC2 modulates T cells in TME by regulating TGFβ-1 expression. The finding that a protein has both tumor intrinsic and extrinsic roles in tumorigenesis is not a novel concept, as illustrated by various groups describing the role of K-ras in regulating immunogenicity in the TME in addition to an oncogenic tumor-intrinsic role (42–45). We found that TGFβ-1 expression was highly elevated in media from PDAC cultures and that VPAC2-deficient PDAC cells had reduced TGFβ-1 secretion, leading to T-cell activation. Transcriptionally, VPAC2 signaling enhances SP1 activity in the nucleus, promoting TGFβ-1 mRNA synthesis, as evidenced by reduced nuclear SP1 levels in VPAC2KO and VPAC2KD cell lines and reduced TGFβ-1 expression after treatment with an SP1 inhibitor. While the transcriptional regulation of TGFβ-1 expression by SP1 has been previously reported in the literature (27–30), this is the first report of SP1 regulation downstream of VPAC2 signaling that can contribute to the malignant progression of PDAC. However, we cannot rule out the other transcription factors profiled from our assay including ELK, OCT4 and p53 as potential regulators downstream of VPAC2 signaling.

The non-cell autonomous effect of VPAC2-expressing tumor cells on the T cells was further supported by *in vivo* data with significantly lower tumor burden in recipients of VPAC2KD subcutaneous and orthotopic KPC.luc tumors. This effect was dependent on the presence of CD4 and CD8 T cells. Thus, our study presents a novel role for VPAC2 receptor on PDAC cells, promoting T cell exclusion and suppression in the TME in a paracrine fashion via TGFβ-1. TGFβ-1 is a pleiotropic cytokine and can have tumor-suppressing or promoting properties during PDAC progression (46–50). TGFβ signaling on cancer cells can induce apoptosis during the premalignant stage, whereas it promotes EMT transition and metastasis of malignant cancer cells (50). However, the role of TGFβ as a negative regulator in the adaptive or innate immune system is well established and thus reflect an important pathway to control the immunogenicity in the TME (47,48,51). While our present study shows modulation of TGFβ-1 secretion by cancer cells via VPAC2 signaling, alternate mechanisms may contribute to better tumor control by T cells in the VPAC2KO/KD tumors. For example, augmented interferon signaling in cancer cells can lead to the expression of inhibitory ligands on cancer cells that can bind to inhibitory receptors on T cells that suppress T cell responses (52–55). To examine this possibility, we interrogated the expression of PDL1 and CD80 following interferon-gamma stimulation. We observed significantly reduced PDL1 induction but increased CD80 expression in the VPAC2KO Panc02 cells versus parental Panc02 cells (**Suppl. Fig.S3E and F**), suggesting VPAC2 signaling may interact with interferon-gamma receptor signaling to enhance immunogenicity in the TME. We previously published that blockade of VIP-receptors using ANT308, a high-affinity VIP-receptor antagonist, enhanced T cell activation and anti-cancer immunity in PDAC (9). Findings from our current study suggest that our previous report showing the anti-cancer activity of the VIP-receptor antagonist in PDAC models could be attributed partially to tumor-cell autonomous and non-tumor-cell autonomous effects from VPAC2 signaling in PDAC cells as defined in this paper. Indeed, treatment of PDAC cell lines with ANT308 resulted in similar effects on colony formation as were seen using the VPAC2 inhibitor (data not shown). In conclusion, our study elucidates the importance of autocrine VIP signaling via VPAC2 as a potential resistance mechanism to promote tumor growth in a tumor cell-intrinsic and extrinsic manner. Within this mechanism, VPAC2 regulates Piwil2 and TGFβ-1 to increase cancer cell growth and T-cell suppression, respectively, leading to reduced tumor control in PDAC (**Fig.7**). Underlying the mechanisms of VIP-receptor signaling in PDAC provides new insights to optimize the clinical development of drugs targeting this highly aggressive cancer.

**Figure 7.**
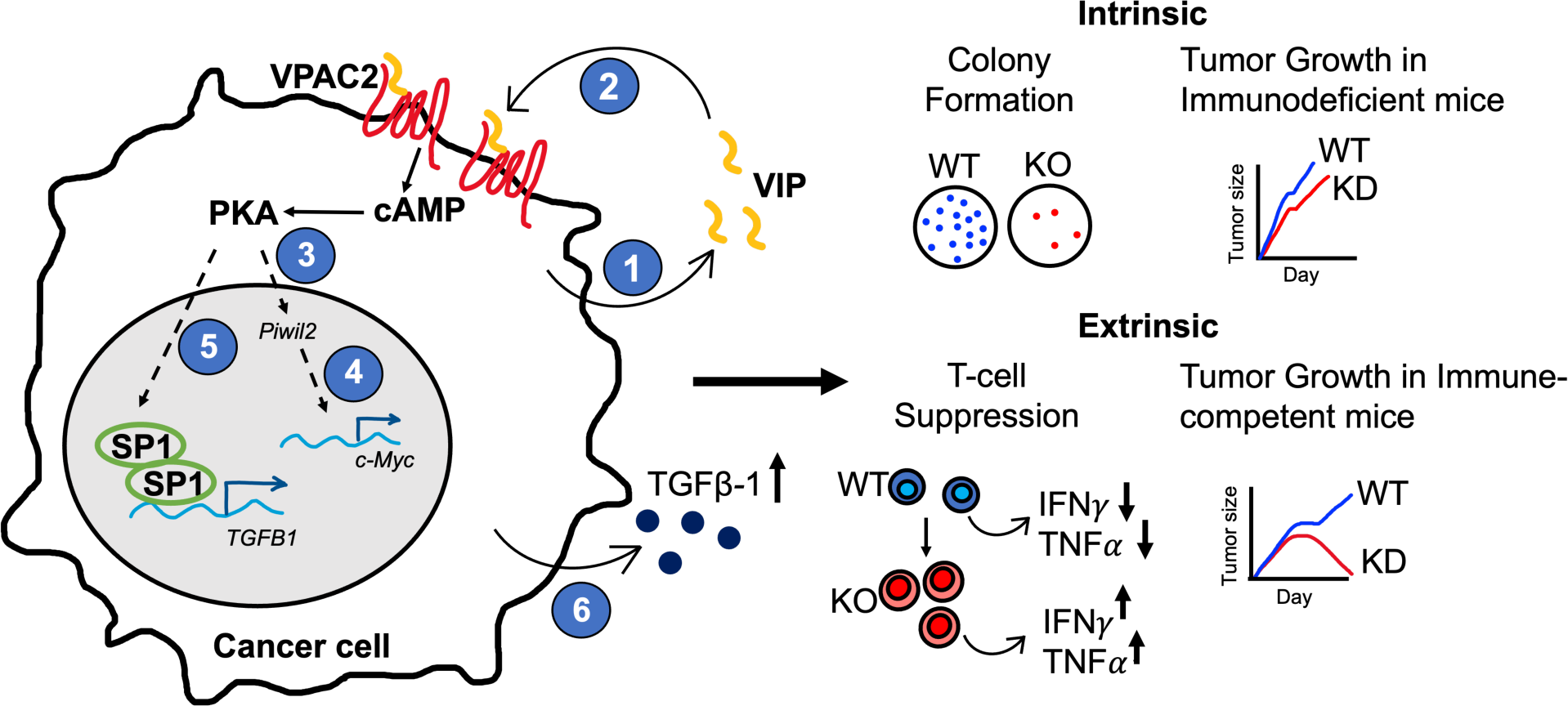
Proposed model showing VPAC2 signaling on PDAC cells driving tumor-intrinsic and tumor-extrinsic effects. 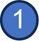 Pancreatic cancer cells overexpress VIP and, 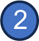 VIP binds to its VIP receptor, VPAC2, in an autocrine manner. Signaling via VPAC2 receptor leads to 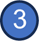 increased Piwil2 expression to promote growth of tumor cells by driving 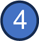 expression of c-myc and its targets. In addition, VPAC2 signaling 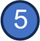 increases active SP1 in the nucleus to drive 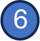 TGFβ-1 expression leading to tumor cell extrinsic effects that suppress T cells as a mechanism of immune escape in the tumor microenvironment. Dotted arrows 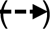 represent undetermined pathways. Representative findings from experiments utilizing wild type (WT, blue) and VPAC2 knock out (KO, red) or knock down (KD, red) are summarized to the right.

## Acknowledgements

The authors thank patients for their tissue samples. The authors also thank Maggie Phillips from Emory University for teaching the procedures of orthotopic surgeries, the shared resources at Emory University, namely Emory Flow Cytometry Core (EFCC), Cancer Animal Models Shared Resource (CAMS), Cancer Tissue Pathology Core (CTP), and Integrated Cellular Imaging Core (ICI), that provided services or instruments at subsidized cost to conduct some of the reported experiments. The authors thank Dr. Tuveson (Cold Spring Harbor Laboratory, Cold Spring Harbor, NY), Dr. Logsdon (MD Anderson Cancer Center, Houston, Texas), and Dr. Pilon-Thomas (H. Lee Moffitt Cancer Center, Tampa, FL) for providing the PDAC cell lines. The data regarding *VIP* or *VIPR1* (encoding VPAC1) and *VIPR2* (encoding VPAC2) mRNA levels in human tumors discussed here are based upon data generated by The Cancer Genome Atlas Research Network: https://www.cancer.gov/tcga. This work was supported in by Katz Foundation funding.

## Author contributions

TP conceptualized the study, designed, and performed experiments, analyzed data, and wrote the manuscript. FZ assisted in the analysis of patient Kaplan-Meir plots. SW, HZ, CH, JML assisted in some *in vivo* experiments and reviewed the manuscript. WW performed *in vitro* experiments and reviewed the manuscript. YL reviewed statistical analyses. SR provided input in experimental design, and GBL provided clinical samples and critically reviewed the manuscript. EKW conceptualized the study, edited the manuscript, and provided funding.

## Notes

**Disclosures of potential conflicts of interest** Intellectual property related to the use of peptide antagonists to vasoactive intestinal polypeptides to treat cancer is the subject of US patent applications with SR, TP, JML, and EKW listed as inventors. These patents have been licensed to Cambium Oncology, LLC. EKW are co-founders and have equity in Cambium Oncology. A conflict-of-interest management plan has been reviewed and approved by Emory University.

### Competing Interest Statement

Intellectual property related to the use of peptide antagonists to vasoactive intestinal polypeptides to treat cancer is the subject of US patent applications with SR, TP, JML, and EKW listed as inventors. These patents have been licensed to Cambium Oncology, LLC. EKW are co-founders and have equity in Cambium Oncology. A conflict-of-interest management plan has been reviewed and approved by Emory University.

