## Supplementary Information for "VPAC2 receptor signaling promotes pancreatic cancer cell growth and decreases the immunogenicity of the tumor microenvironment"

### Supplementary Materials and Methods

*Western Blot (WB)*

Cells were lysed on ice for 20 minutes with 1X RIPA Lysis buffer (Thermo Scientific) supplemented with protease inhibitor cocktail and phosphatase inhibitor cocktail. Cell homogenate was vortexed every 5 minutes. Supernatant containing cell lysate was collected by spinning down at 15,000 rpm at 4ºC for 20 minutes. Antibodies used for WB are detailed in Supplementary Table 1.

*Flow cytometry*

Single cell suspensions from spleen and tumor harvested from KPC.luc bearing mice were prepared as described before (1). Single cells were stained with Fixable Aqua live/dead stain followed by Fc block for 5 minutes each. Surface antibodies were added to the cells at the desired concentration and left to stain for 30 minutes at 4ºC. Cells were fixed and permeabilized for intracellular staining according to manufacturer's protocol (Cat. 00-5523-00, Invitrogen™). For CFSE staining, cells were stained with CFSE dye (Cat. C34554, Invitrogen™) at 0.5μM for 15 minutes at 37°C and used in subsequent studies.

**Supplementary Table 1.** Details of antibodies used in this study. Abbreviations; WB for Western Blot, IF for Immunofluorescence, IHC for Immunohistochemistry, FC for flow cytometry)

| Target species | Target | Clone | Vender | Catalog no. | Dilution |
|---|---|---|---|---|---|
| ***In vivo* MAb** | PD-1 | RMP1-14 | BioXcell | BE0146 | 200μg (X4) |
| | Rat IgG2A Isotype control | 2A3 | BioXcell | BE0089 | 200μg (X4) |
| | CD4 | GK1.5 | BioXcell | BE0003-1 | |
| | CD8 | Clone 2.43 | BioXcell | BE0061 | |
| **Other Antibodies** | P-CREB (S133) | 87G3 | Cell Signaling | 14001S | 1:50 (FC) |
| | SP1 | 9389S | Cell Signaling | 9389S | 1:1000 (WB) |
| | VPAC1 | SP234 | Abcam | ab183312 | 1:500 (WB) |
| | VPAC2 | | Invitrogen | PA5-21304 | 1:500 (WB) and 1:10 (FC) |
| | | SP235 | Abcam | ab183334 | 1:200 (IF) |

| | | | | | |
|---|---|---|---|---|---|
| | VIP | OTI5B5 | Origene | CF806852 | 1:50 (IF) |
| | | | Genetex | GTX129461 | 1:500 |
| | Cytokeratin 19 (CK19) | EP1580Y | Abcam | ab52625 | 1:400 (IF) |
| | | | Invitrogen | PA5-143131 | 1:500 (IF) |
| | CD4 | EPR6855 | Abcam | ab133616 | 1:500 (IHC) |
| | CD8 | EP1150Y | Abcam | ab93278 | 1:500 (IHC) |
| | Ki67 | SP6 | Abcam | ab16667 | 1:200 (IHC) |

**Supplementary Table 2.** The List of primers used for PCR or qPCR.

| **Target** | **Forward Primer (5'-3')** | **Reverse Primer (5'-3')** |
|---|---|---|
| ***18S*** | CGGACAGGATTGACAGATTGATAGC | TGCCAGAGTCTCGTTCGTTATCG |
| ***GAPDH-1*** | AGGAGAGTGTTTCCTCGTCCC | CAGATCCACGACGGACACAT |
| ***GAPDH-2*** | AACGACCCCTTCATTGAC | TCCACGACATACTCAGCAC |
| ***VIP (exon5)*** | TCACTGACAACTATACCCGCC | CCTCACTGCTCCTCTTTCCA |
| ***VIPR2 (exon2)*** | GCATCCATCCAGAATGTCGC | GCCAGCATGTGATGTTGTCC |
| ***VIPR2 (exon9-12)*** | ATGGACAGCAACTCGCCTCTCTTTAG | GGAAGGAACCAACACATAACTCAAACAG |
| ***Piwil2*** | TGTCACATCGGAGGACCTGAAC | CCAGTTCTCTGGCTTGGTCCAT |

**Supplementary Figures**

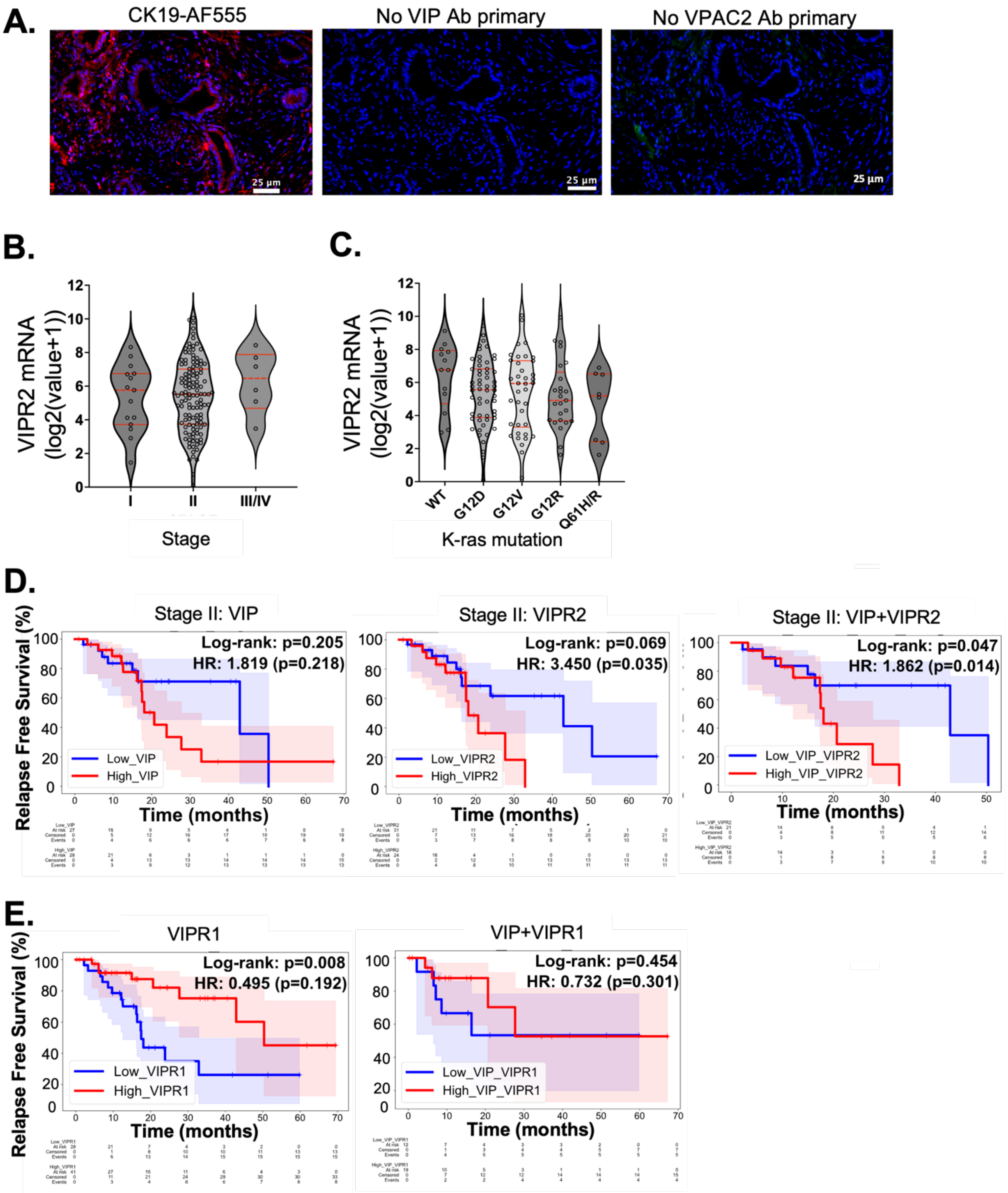


**Supplementary Figure S1**. High VIP and VPAC2 but not VPAC1 expression in PDAC patients correlate with worse relapse free outcomes. **(A)** Immunofluorescence staining control for VIP and VPAC2 in human PDAC tissue using only secondary antibodies conjugated to fluorochromes. Cytokeratin 19 (CK19) staining shown for pancreatic cancer ductal cells. **(B)** VPAC2 mRNA expression in PDAC patients stratified based on stage and **(C)** K-ras mutation status. **(D)** Relapse free survival for stage II PDAC patients with high and low expression of VIP and VPAC2. The shaded colors below and above the survival curves correspond to 95% confidence interval for the respective patient groups. **(E)** Relapse free survival for PDAC patients with high and low expression VPAC1 and high and low expression of VIP and VPAC1. Log rank test was performed for survival difference between patients. Hazard ratio (HR) was estimated by Cox proportional hazard model.

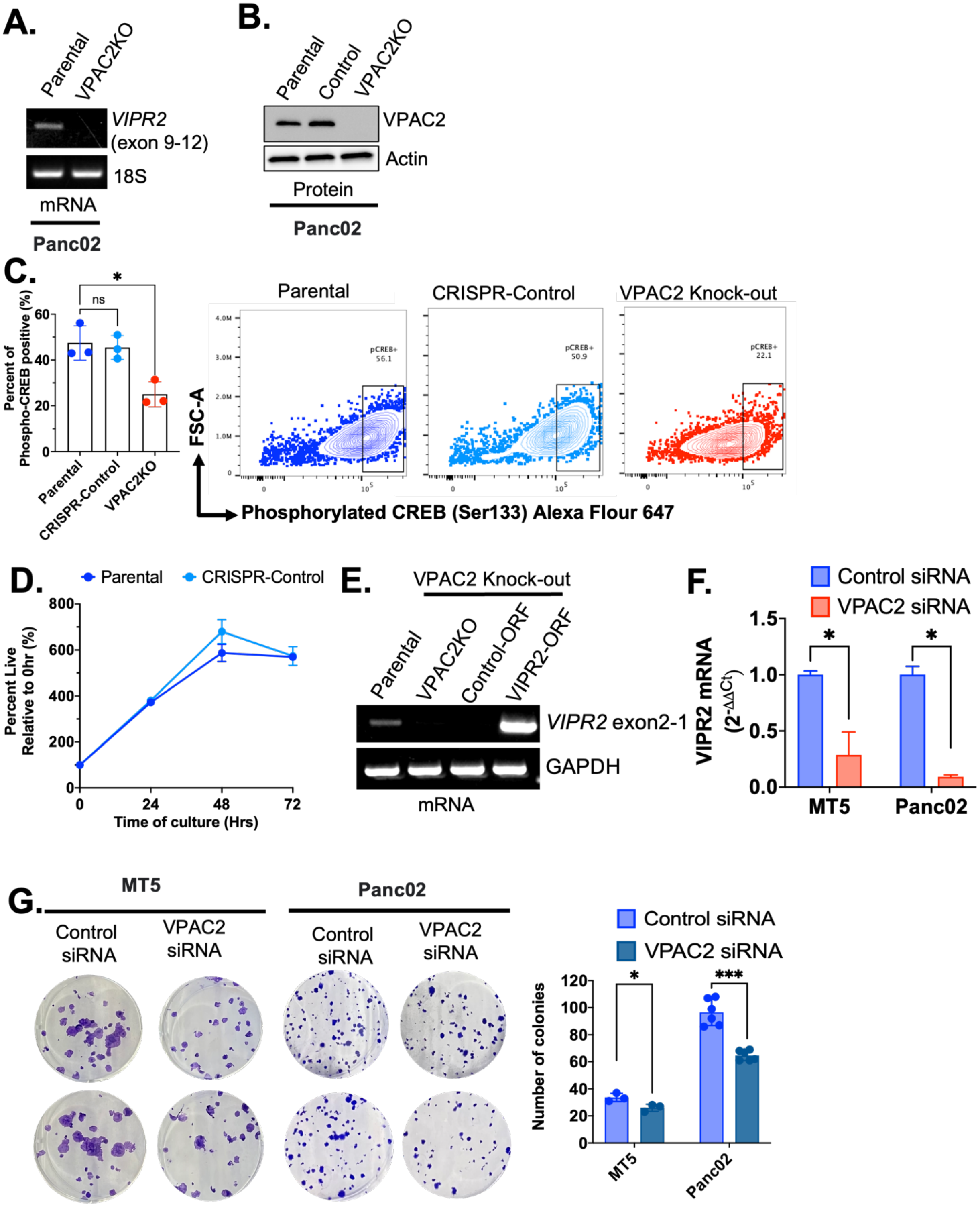


**Supplementary Figure S2**. Absence of VIP and VPAC2 signaling leads to decreased colony formation in vitro. **(A)** RT-PCR and **(B)** Western blot confirmation for VPAC2KO

Panc02 cells. Another clone which underwent CRISPR editing, showing no loss of VPAC2 protein expression and no decrease in **(C)** phosphorylation of CREB is thereby labelled as CRISPR-Control. **(D)** MTT assay showing growth of Panc02 versus CRISPR-Control cells over 72 hours. **(E)** RT-PCR confirmation of VPAC2 rescue (VIPR2-ORF) on VPAC2KO **(F)** qRT-PCR confirmation of VPAC2 mRNA expression following transfection with control or VPAC2-targeting siRNA in MT5 and Panc02 (n=3). **(G)** Crystal violet colony assay for MT5 and Panc02 cells. siRNA transfected cells were plated at 50-100 cells per 6-well plate and cultured for 10 days. Cells were stained with crystal violet stain after 10 days of culture and number of colonies were computed using countPHICS software. Data are presented as bar graphs or line plots ± standard deviation (SD). For C, one-way ANOVA test following by Dunnet's multiple comparison post-hoc test was performed. For F and G, two-tailed unpaired t test was used. *$p<0.05$, ***$p<0.001$, ****$p<0.0001$.

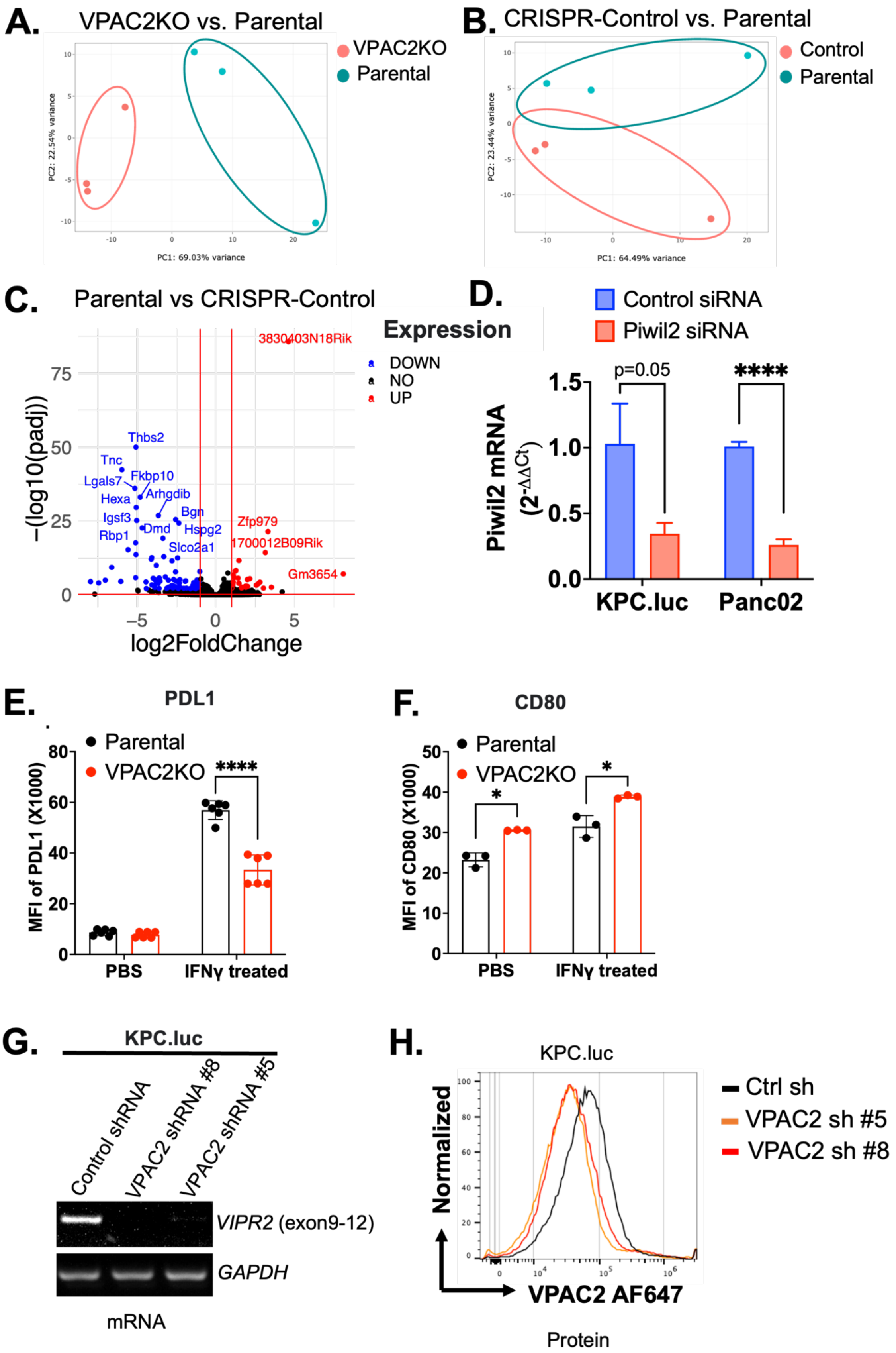
A.
VPAC2KO vs. Parental
VPAC2KO
Parental
PC2: 22.54% variance
PC1: 69.03% variance
B.
CRISPR-Control vs. Parental
Control
Parental
PC2: 23.44% variance
PC1: 64.49% variance
C.
Parental vs CRISPR-Control
3830403N18Rik
DOWN
NO
UP
Thbs2
Tnc
Lgals7
Fkbp10
Hexa
Arhgdib
Igsf3
Bgn
Dmd
Hspg2
Rbp1
Slco2a1
Zfp979
1700012B09Rik
Gm3654
−(log10(padj))
log2FoldChange
D.
Expression
Control siRNA
Piwil2 siRNA
p=0.05
****
Piwil2 mRNA
(2^-ΔΔCt)
KPC.luc
Panc02
E.
PDL1
Parental
VPAC2KO
****
MFI of PDL1 (X1000)
PBS
IFNγ treated
F.
CD80
Parental
VPAC2KO
*
*
MFI of CD80 (X1000)
PBS
IFNγ treated
G.
KPC.luc
Control shRNA
VPAC2 shRNA #8
VPAC2 shRNA #5
VIPR2 (exon9-12)
GAPDH
mRNA
H.
KPC.luc
Ctrl sh
VPAC2 sh #5
VPAC2 sh #8
Normalized
VPAC2 AF647
Protein

**Supplementary Figure S3**. Principal component analysis (PCA) between **(A)** Parental and VPAC2KO **(B)** Parental and CRISPR-Control**. (C)** Volcano plot of significantly downregulated (blue) and upregulated (red) genes between parental and CRISPR-Control Panc02 cells from RNA Sequencing analysis. A total of 145 differentially expressed genes were detected and were eliminated from analysis comparing parental and VPAC2KO. **(D)** qRT-PCR confirmation of Piwil2 mRNA expression in siRNA transfected KPC.luc and Panc02 cells (n=3). **(E)** PDL1 and **(F)** CD80 expression as detected from flow cytometry before and after stimulation with IFN$\gamma$ (10ng/ml) in parental versus VPAC2KO Panc02 cells. **(G)** RT-PCR and **(H)** Flow cytometry confirmation for knock down of VPAC2 in 2 clones (#8 and #5) of KPC.luc cells. For D, data are presented as bar graphs + standard deviation (SD). For E and F, data are presented as ± SD. For D-F, two-tailed unpaired t test was used. **p<0.01, ***p<0.001, ****p<0.0001.

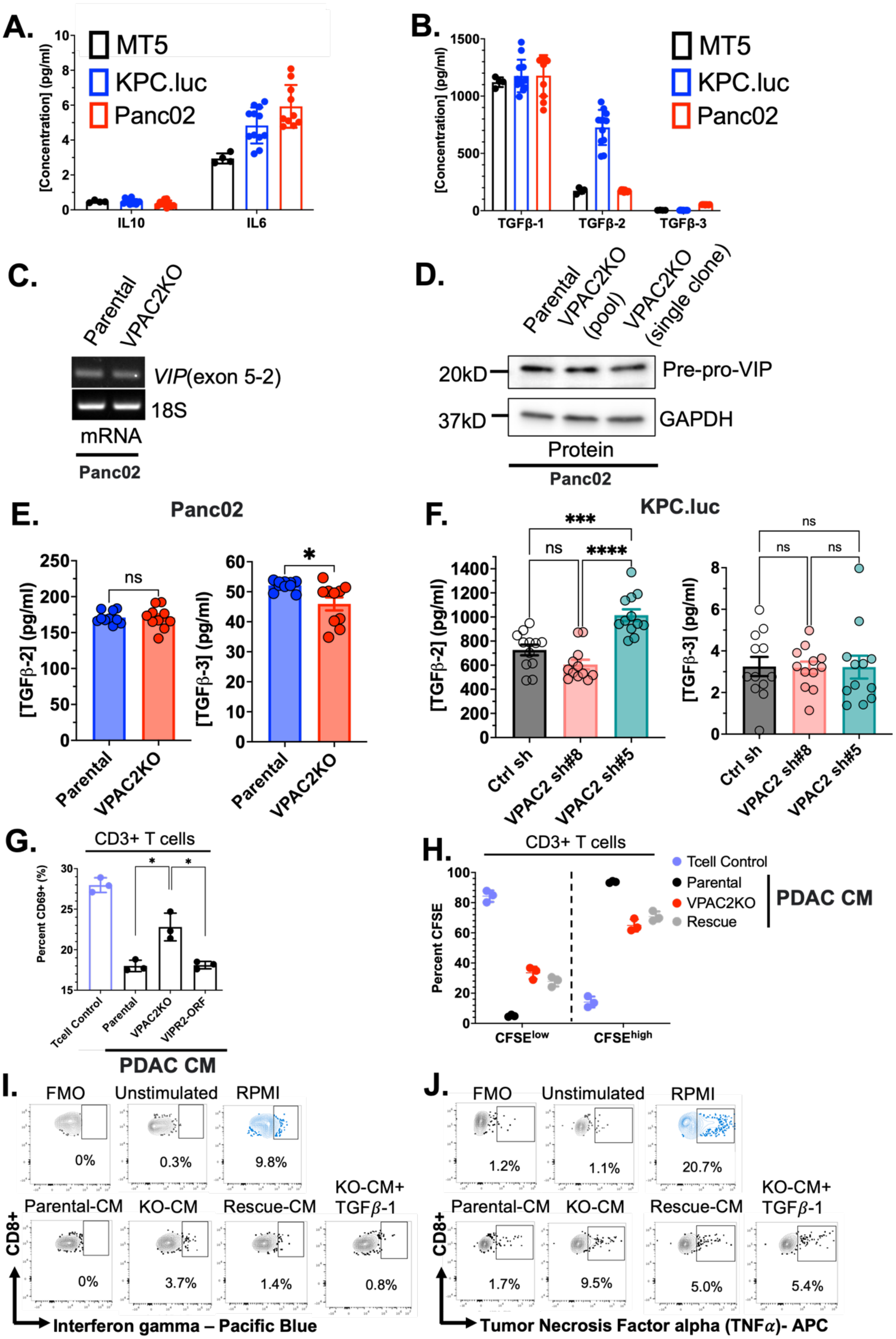

A.
[Concentration] (pg/ml)
MT5
KPC.luc
Panc02
IL10
IL6
B.
[Concentration] (pg/ml)
TGFβ-1
TGFβ-2
TGFβ-3
C.
Parental
VPAC2KO
VIP(exon 5-2)
18S
mRNA
Panc02
D.
Parental
VPAC2KO (pool)
VPAC2KO (single clone)
20kD
Pre-pro-VIP
37kD
GAPDH
Protein
Panc02
E.
Panc02
[TGFβ-2] (pg/ml)
[TGFβ-3] (pg/ml)
ns
*
Parental
VPAC2KO
F.
KPC.luc
***
ns
****
[TGFβ-2] (pg/ml)
[TGFβ-3] (pg/ml)
Ctrl sh
VPAC2 sh#8
VPAC2 sh#5
G.
CD3+ T cells
Percent CD69+ (%)
Tcell Control
Parental
VPAC2KO
VIPR2-ORF
PDAC CM
H.
CD3+ T cells
Tcell Control
Parental
VPAC2KO
Rescue
PDAC CM
Percent CFSE
CFSE^low
CFSE^high
I.
FMO
Unstimulated
RPMI
0%
0.3%
9.8%
Parental-CM
KO-CM
Rescue-CM
KO-CM+ TGFβ-1
0%
3.7%
1.4%
0.8%
CD8+
Interferon gamma – Pacific Blue
J.
FMO
Unstimulated
RPMI
1.2%
1.1%
20.7%
Parental-CM
KO-CM
Rescue-CM
KO-CM+ TGFβ-1
1.7%
9.5%
5.0%
5.4%
CD8+
Tumor Necrosis Factor alpha (TNFα)- APC

**Supplementary Figure S4**. Disruption of VPAC2 pathway does not influence VIP or other isoforms of TGFβ. Levels of secreted cytokines including **(A)** IL10 and IL6 and **(B)** TGFβ-1, TGFβ-2, and TGFβ-3 in MT5, KPC.luc and Panc02. **(C)** mRNA and **(D)** protein expression of VIP in parental versus VPAC2KO Panc02 cells. Levels of secreted TGFβ-2 and TGFβ-3 in **(E)** Panc02 and **(F)** KPC.luc cultures. CRISPR-KO or lentiviral knock down of VPAC2 cells were compared to control cultures. **(G)** Percent of CD69 and **(F)** $CFSE^{low}$ and $CFSE^{high}$ of CD3+ T cells following culture in conditioned media from parental, VPAC2KO, VPAC2-rescue Panc02 cells. T cells were stimulated with 0.1μg/ml anti-CD3 coated plates and cultured for 24 hours and analyzed for CD69 expression by flow cytometry. For CFSE analysis, T cells were cultured for 6 days with 0.1μg/ml anti-CD3 coated plates. Representative flow plots with appropriate gating for **(I)** IFN$\gamma$ positive and **(J)** TNF$\alpha$ positive CD8+ T cells are shown. For A, B, E, and F, data are presented as bar graphs ± standard error (SEM). For G and H, data are presented as ± standard deviation (SD). For E, F and G, unpaired t test with Welch's correction was used. *p<0.05, ***p<0.001, ****p<0.0001.

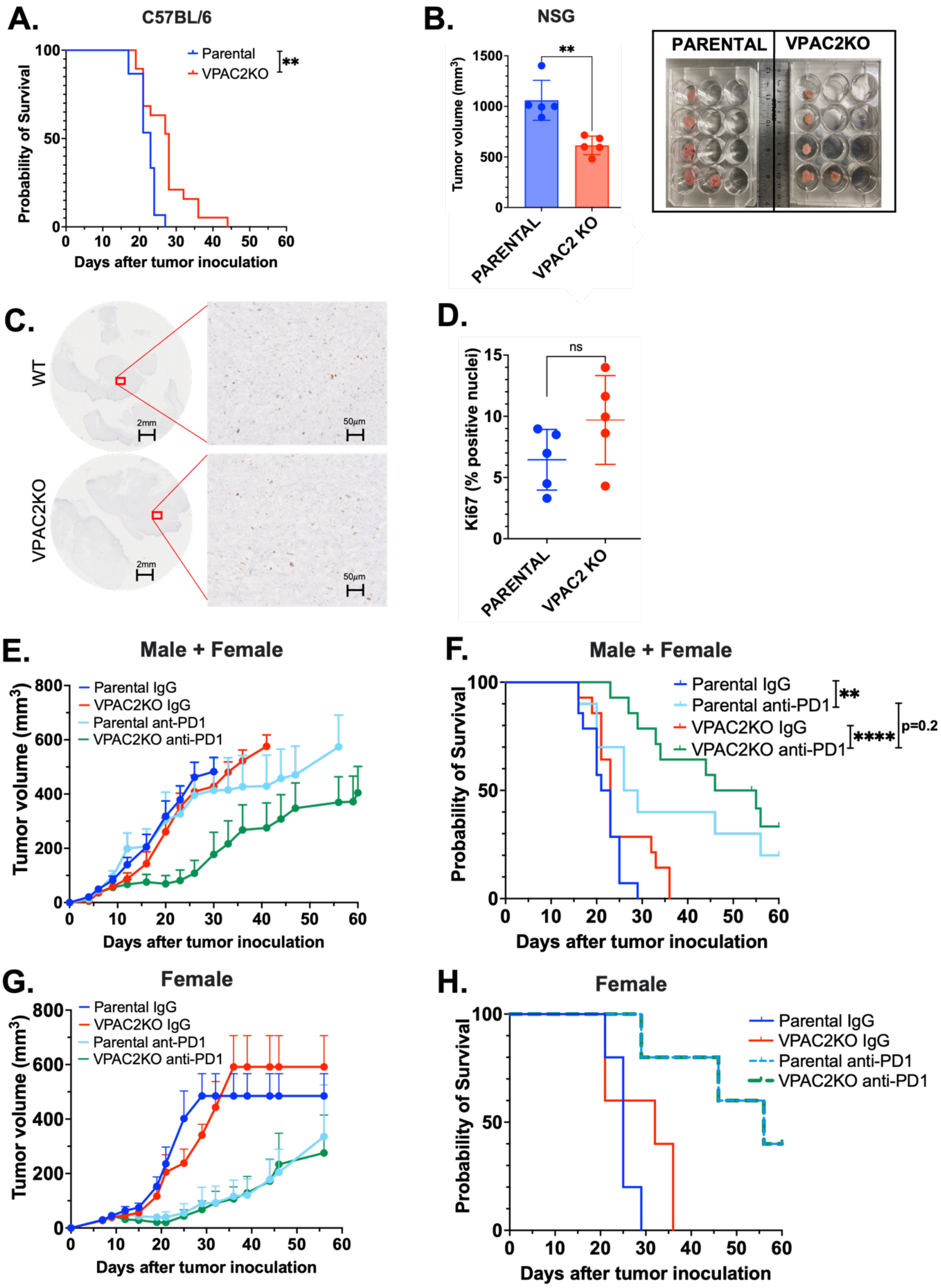
A.
C57BL/6
Parental
VPAC2KO
**
Probability of Survival
Days after tumor inoculation
B.
NSG
Tumor volume (mm³)
**
PARENTAL
VPAC2 KO
PARENTAL
VPAC2KO
C.
WT
VPAC2KO
2mm
50µm
D.
Ki67 (% positive nuclei)
ns
PARENTAL
VPAC2 KO
E.
Male + Female
Parental IgG
VPAC2KO IgG
Parental ant-PD1
VPAC2KO anti-PD1
Tumor volume (mm³)
Days after tumor inoculation
F.
Male + Female
Parental IgG
Parental anti-PD1
VPAC2KO IgG
VPAC2KO anti-PD1
**
****
p=0.2
Probability of Survival
Days after tumor inoculation
G.
Female
Parental IgG
VPAC2KO IgG
Parental ant-PD1
VPAC2KO anti-PD1
Tumor volume (mm³)
Days after tumor inoculation
H.
Female
Parental IgG
VPAC2KO IgG
Parental anti-PD1
VPAC2KO anti-PD1
Probability of Survival
Days after tumor inoculation

**Supplementary Figure S5**. Absence of VPAC2 leads to decreased tumor growth *in vivo* in a tumor intrinsic and extrinsic manner in subcutaneous Panc02 model. **(A)** Kaplan-Meir survival plots of C57BL/6 mice implanted with Parental versus VPAC2KO Panc02 cells (n=15-20). **(B)** Tumor volume at day 21 post tumor implantation in NOD SCID gamma (NSG) mice (n=5). **(C)** Representative images for **i**mmunohistochemistry for Ki67 on parental and VPAC2KO tumors in NSG mice at day 21. **(D)** Number of Ki67 positive nuclei were computed using QuPath software. **(E)** Combined male and female data for average tumor volume over time for parental and VPAC2KO Panc02 cells injected to C57BL/6 (n=10-15) and treated with isotype IgG or anti-PD1 antibody. **(F)** Kaplan-Meir survival plots corresponding to E. Tumor volumes were measured by vernier calipers one to two times a week until study endpoint. **(G)** Average tumor volume over time for parental and VPAC2KO Panc02 cells injected to female C57BL/6 (n=5) and treated with isotype IgG or anti-PD1 antibody. **(H)** Kaplan-Meir survival plots corresponding to G**.** For B and D, data are presented ± standard deviation (SD) and two-tailed unpaired t test was used. All other data are presented + standard error (SEM). Log-rank test was used for statistical differences for Kaplan-Meier curves. **p<0.01 and ****p<0.0001.

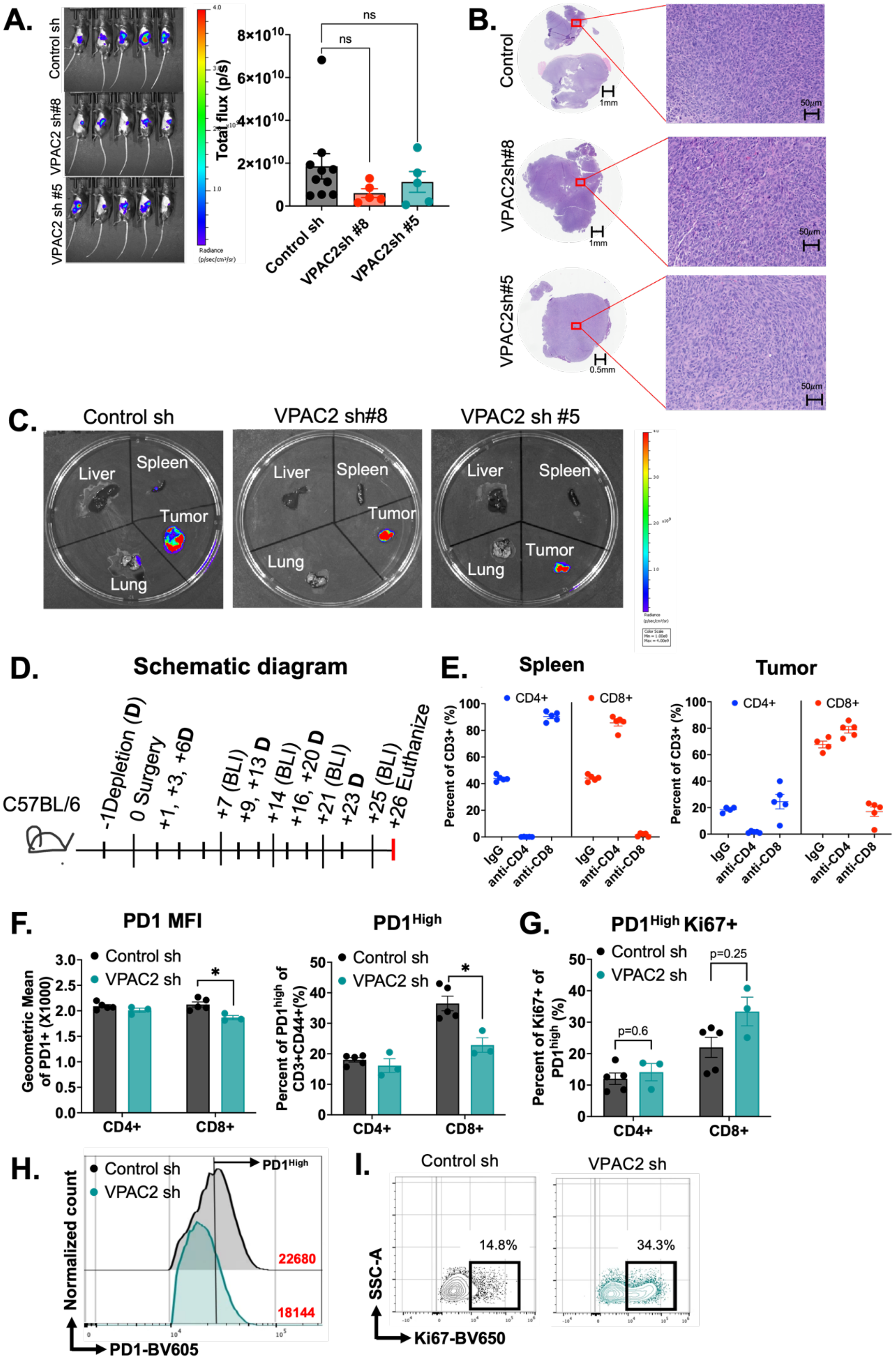
A.
Control sh
VPAC2 sh#8
VPAC2 sh #5
Total flux (p/s)
8×10^10
6×10^10
4×10^10
2×10^10
0
ns
ns
Control sh
VPAC2sh #8
VPAC2sh #5
B.
Control
VPAC2sh#8
VPAC2sh#5
1mm
0.5mm
50μm
C.
Control sh
VPAC2 sh#8
VPAC2 sh #5
Liver
Spleen
Tumor
Lung
D.
Schematic diagram
C57BL/6
-1Depletion (D)
0 Surgery
+1, +3, +6D
+7 (BLI)
+9, +13 D
+14 (BLI)
+16, +20 D
+21 (BLI)
+23 D
+25 (BLI)
+26 Euthanize
E.
Spleen
Tumor
CD4+
CD8+
Percent of CD3+ (%)
IgG
anti-CD4
anti-CD8
F.
PD1 MFI
Control sh
VPAC2 sh
Geometric Mean of PD1+ (X1000)
CD4+
CD8+
PD1High
Percent of PD1high of CD3+CD44+(%)
G.
PD1High Ki67+
Percent of Ki67+ of PD1high (%)
p=0.6
p=0.25
H.
Normalized count
PD1High
22680
18144
PD1-BV605
I.
Control sh
VPAC2 sh
14.8%
34.3%
SSC-A
Ki67-BV650

**Supplementary Figure S6**. Knock down of VPAC2 on does not influence metastatic potential in KPC.luc orthotopic model. Control sh and VPAC2 sh (#8 and #5) KPC.luc cells were surgically implanted in the tail of the pancreas of C57BL/6 mice. **(A)** IVIS bioluminescent image and total flux measured at day 7 post implantation. Isoflurane was used for anesthesia for bioluminescent imaging. **(B)** Representative images from H&E stained orthotopic tumors at day 26 when mice were euthanized. **(C)** Representative pictures from IVIS bioluminescent imaging acquired at day 26 of liver and lung relative to tumor showing no gross metastasis at the respective organs. Spleen shown as negative control. **(D)** Schematic diagram showing orthotopic implantation of control and VPAC2 sh KPC.luc cells and depletion strategy for anti-CD4 and anti-CD8 blockade until study end point. **(E)** Scatter dot plots showing percent CD4+ and CD8+ T cells in spleen and tumor with or without anti-CD4/anti-CD8 blockade. Following euthanasia at day 26, KPC.luc tumors were dissociated as single cells and analyzed for T cell phenotype by flow cytometry. Bar graph showing **(F)** Geometric mean fluorescent intensity (MFI) of PD1 positive (left) and percent of T-cells expressing $PD1^{High}$ (right) and **(G)** $PD1^{High}$Ki67 positive as gated on CD4+ or CD8+. **(H)** Representative histogram for PD1 positive gated on CD8+CD44+ for F. The numbers in red represent the geometric mean value for the histogram. The arrow from the vertical line indicates the gating for $PD1^{High}$ cells for F. **(I)** Representative contour plots for Ki67 positive cells as gated on $PD1^{High}$ cells for G. All data are presented as ± standard error (SEM). For A, one-way ANOVA test following by Dunnet's multiple comparison post-hoc test was used. For F and G, Mann-Whitney test was used. *$p<0.05$
